# Maternal high-fat diet drives sex-specific microglia remodeling of serotonergic reward circuitry

**DOI:** 10.64898/2026.05.10.723768

**Authors:** Michael S. Patton, William Sun, Lillian Stanley, Aasha Paredes, Joanna Kang, Zachary Schettewi, Benjamin Horvath, Julia E. Dziabis, Benjamin A Devlin, Trisha V. Vaidyanathan, Staci D. Bilbo

## Abstract

Maternal nutrition shapes offspring brain development and influences neurodevelopmental disorder risk, but the underlying mechanisms remain unclear. In mice, maternal high-fat diet exposure disrupted microglia-serotonin interactions during a critical postnatal period, producing persistent, sex-specific mesolimbic alterations. Male but not female offspring showed increased serotonergic fiber density in the nucleus accumbens (NAc), coincident with reduced microglial phagocytosis of serotonergic projections. Microglial 5-HT2C receptor signaling is a key regulator of this process. Viral over expression in microglia, mimicking diet-induced upregulation, was sufficient to cause serotonergic hyperinnervation. By adulthood, male offspring displayed increased NAc serotonin release and projection-specific changes in dorsal raphe physiology. These circuit alterations accelerated reward-motivated learning, a phenotype reproduced by chemogenetic activation of NAc-projecting serotonergic neurons. Together, these findings reveal a novel mechanism by which maternal diet programs serotonergic circuit assembly and behavior in a sex-specific manner, providing a potential link between early-life metabolic inflammation and lifelong serotonergic dysfunction.

## Introduction

Maternal nutrition is a powerful regulator of fetal development and exerts long-lasting effects on offspring physiology and disease vulnerability^1–3^. Diets high in saturated fats contribute substantially to excess gestational weight gain^4, 5^, which is epidemiologically associated with higher incidences of neurodevelopmental disorders in children^6–10^. Experimental studies in rodents^11–14^ and non-human primates^15–17^ corroborate these findings, demonstrating that maternal high-fat diet (mHFD) exposure impairs offspring cardiovascular and neural outcomes. Mechanistically, mHFD induces a systemic proinflammatory state^18–22^, characterized by elevated cytokines (e.g.,TNF-a, IL-6, IL-33) that activate immune cells in the developing fetal brain^23, 24^, perturbing neuroimmune interactions that orchestrates brain development^25–28^. Despite these advances, it is still unknown whether maternal metabolic state broadly impacts neural circuit development, or whether specific circuit vulnerabilities underly disease pathology, warranting more targeted therapeutic interventions.

Microglia, the resident immune cells of the central nervous system, are central to brain development, regulating neural progenitors^29, 30^ and sculping emerging circuits through axon guidance^31–34^ and synaptic pruning^35–38^. Microglia integrate both neural and immune cues, making them ideally positioned to sense maternal inflammatory signals elicited by mHFD. Consistent with the maternal immune activation framework, transient immune perturbation during sensitive developmental windows can reprogram microglial state, disrupting microglia-neuron interactions required for proper circuit assembly. Such early immune programming produces enduring alterations in neural connectivity and function that are thought to underlie the behavioral abnormalities observed in adult offspring^39–43^. We recently demonstrated that mHFD increases microglia phagocytosis of serotonin in the male embryonic day 14.5 dorsal raphe nucleus (DRN)^11^, the principal source of forebrain serotonin^44^. Consistent with this finding, mHFD exposure in macaques reduces the density of DRN serotonergic neurons^45^. Together, these results identify the developing serotonergic system as a target of mHFD-induced inflammation and suggests a mechanistic link between early immune perturbations and the serotonergic dysfunction commonly observed across neurodevelopmental disorders^46–50^.

The serotonin system is among the earliest neuronal networks to emerge, appearing in the mouse brain by embryonic day 11.5^51^. By late gestation, serotonergic axons reach their target fields^51, 52^ and continue to mature throughout the first three postnatal weeks^53^. This extended developmental window renders the serotonin system particularly sensitive to early-life perturbations, as serotonergic axons demonstrate the remarkable capacity to modify axonal branching in response to early-life challenges^54^. Here we investigated how mHFD influences postnatal development of serotonin circuits and found that mHFD selectively increases serotonergic fiber innervation to the nucleus accumbens (NAc) of male but not female postnatal (P14) offspring. This hyperinnervation coincides with reduced microglial phagocytosis of serotonergic fibers in the NAc. Using a novel viral approach to manipulate microglia, we show that over expressing the 5-HT 2C receptor, mimicking its upregulation in NAc microglia following mHFD, induces hyperinnervation of serotonergic fibers in postnatal (P14) offspring. By adulthood, mHFD male offspring exhibit elevated serotonergic innervation to the NAc, greater serotonin release, and projection-defined alterations in DRN neuronal physiology. Finally, we show that mHFD male offspring exhibit accelerated reward-motivated learning, a phenotype recapitulated by chemogenetic activation of NAc-projecting serotonergic neurons.

## Methods

### Animals

All procedures relating to animal care and treatment conformed to Duke Institutional Animal Care and Use Committee (IACUC) and NIH guidelines (protocol number A062-22-03). Animals were group housed in a standard 12:12 light-dark cycle (lights off at 0900h, on at 2100h) with *ad libitum* access to food and water unless otherwise specified (see voluntary ethanol consumption, circadian behavior, and CPP). The following mouse lines were used in this study: *C57BL/6J* (JAX #000664), *SERT-cre* (JAX #014554), SELECTIV mice (JAX #037553),GCaMP8s mice (JAX #037952), Cx3cr1-GFP (JAX #005582), RiboTag mice (JAX #029977), and Cx3cr1-Cre BAC-transgenic^55^. The Cre transgene was maintained in males for all experimental studies.

### Diets

4-weeks old female *C57BL/6J* mice were randomly assigned to either a high-fat diet (45% kcal from fat; Research Diets D12451i) or low-fat diet (10% kcal from fat; Research Diets D12450Hi). For more information regarding diet composition, see Tables 1-3. After 6 weeks on diet, females were mated with males and dams were maintained on their assigned diet chow throughout gestation and until the pups were weaned (P28). At weaning, offspring were group housed with sex-matched littermates and provided *ad libitum* access to rodent chow 5001 (13% kcal from fat; standard rodent chow Lab Diet 5001).

### Immunohistochemistry

Mice were euthanized with CO_2_ and transcardially perfused with saline followed by ice-cold paraformaldehyde (4%). Brains were extracted and post-fixed in paraformaldehyde (4%) for 24-48h at 4°C. Fixed brains were cryoprotected in 30% sucrose + 0.1% sodium azide in PBS for 3 days or until brains sunk and then flash frozen on dry ice and stored at −80°C until further processing. Frozen brains were cryo-sectioned using a Leica cryostat (CM1950). Sections were blocked at room temperature (RT) with 10% goat serum in PBS + 0.3% Triton X-100 for 2-4h. Sections then incubated in primary antibody overnight at RT. For antibody information, see Table 4. Sections were then washed in PBS and incubated in secondary antibodies for 2-4h at RT. Sections were mounted and coverslipped with Fluoromount-G mounting medium with DAPI (ThermoFisher #00-4959-52). To visualize biotin filled neurons, sections were washed in 0.1 M PBS (3 x 5 min) and blocked with 1% BSA in PBS + 0.3% Triton X-100 (PBS-T) for 2-4 h at room temperature. Slices were incubated with streptavidin Alexa-647 conjugated probe (1:1000, 1% BSA in 0.3% PBS-T; Invitrogen #S21374) for 1h at room temperature.

### RNA-FISH

RNA-FISH was conducted following the manufacturer’s protocol (Molecular Instruments). For DRN analysis, 45 μm sections from virally transfected SERT-cre mice incubated overnight at 37°C in primary probe solutions targeting *Gad2* (1:50 dilution), *Vglut3* (1:50 dilution), and *mCherry* (1:50 dilution). For NAc analysis, 45 μm sections from virally transfected SELECTIV:Cx3CR1-BAC-cre mice incubated overnight at 37°C in IBA1 antibody (1:500 dilution). Excess primary was removed the following day with PBS-T washes. Sections then incuabed in secondary antibody for 4h at room temperature. Sections were washed with 5X SSCT and post-fixed for 10min at room temp with 4% PFA. Sections were then incubated in primary probe solution targeting *Htr2c* overnight at 37°C. Excess probes were removed on the following day with successive washes of increasing concentrations of 5X SSCT (25%, 50%, 75%, 100%) at 37°C. To enable RNA detection via fluorescence, tissue sections were incubated with hairpin probes containing fluorophores. Sections were incubated with 6 pmol of hairpin H1 and 6 pmol of hairpin H2 for a final concentration of 20nM per hairpin in 200μL of the manufacturer’s amplification buffer. Sections were then washed with 100% 5X SSCT, mounted, and coverslipped using Fluoromount-G mounting medium with DAPI.

### Confocal microscopy

All images were collected using an Olympus Fluoview confocal microscope (FV3000). For microglia reconstructions and serotonin fiber density analysis images were collected using a 60x NA1.3 objective (Olympus) with a 2x digital zoom, 20μm z-depth, z-step 0.45μm. For dye-filled neuronal Sholl analysis, images were collected using a 30x 1.05NA objective (Olympus), 80-120μm z-depth, 1μm z-step. For adult synapse analysis, images were collected on a 60x NA1.3 objective with 2.13 digital zoom, 10μm z-depth, 0.45μm z-step. For DRN projection analysis, images were collected using a 10x NA 0.3 objective (Olympus), 10-30μm z-depth, 15μm z-step. For CTB analysis, images were collected using a 20x 0.8NA objective (Olympus), 10-15μm z-depth, 0.9μm z-step. For DRN RNA-FISH, images were collected using a 30x objective, 10μm z-depth, 0.75μm z-step. For microglia RNA-FISH, images were collected using a 60x objective, 2x digital zoom, 20μm z-depth, 0.5μm z-step. For viral histology confirmation, images were collected using a 60x NA1.3 objective with a 1.26 digital zoom, 15-30μm z-depth, 0.9μm z-step.

### IMARIS

3D reconstructions of microglia were conducted using the surface rendering function in IMARIS imaging software (version 10). Lysosomal capacity was calculated by reconstructing individual microglia volume, then masking CD68 fluorescent signal within individual cells. This CD68 signal was then reconstructed and the volume of CD68 was calculated as a percent of the total microglia volume. SERT was similarly reconstructed within CD68 and the volume and calculated as the percent of the total microglia volume.

### Surgical procedures

Adult stereotaxic injections were performed under isoflurane anesthesia (5% induction; 2%–3% maintenance) and viruses were injected at a rate of 40nl/min with a 25G syringe (Hamilton Company). To isolate dorsal raphe nucleus (DRN) serotonergic neurons projecting to the orbitofrontal cortex (OFC) or nucleus accumbens (NAc), SERT-cre mice were bilaterally injected with a retrograde virus expressing flip recombinase cre-dependently (AAVrg pEF1a-DIO-FLPo-WPRE-hGhpA; Addgene #87306) in the OFC (at 10° angle: AP +2.3 mm, ML ± 1.85 mm, DV −2.0 mm; 300 nl/hemisphere) or the NAc (at 10° angle: AP +1 mm, ML ± 1.7 mm, DV −4.0mm; 400 nl/hemisphere). For patch-clamp electrophysiology and viral 3D mapping, SERT-cre mice were also injected with a virus expressing a flip-dependent mCherry (AAV8 Ef1a fDIO-mCherry; Addgene #114471) in the DRN (at 20° angle: AP −4.35 mm, ML ± 0.8 mm, DV −2.6mm; 300 nl). For the DREADDs study, SER-cre mice were injected with the same rgAAV in the NAc and the excitatory DREADD (AAV8 hSyn fDIO-hM3D-mCherry-WPREPa; Addgene #154868), the inhibitory DREADD (AAV8 hSyn fDIO-hM4Di-mCherry-WPREPa; Brain VTA), or control fluorophore (AAV8 hSyn fDIO-mCherry-WPRE-hGHpA; BrainVTA) in the DRN. For serotonin projection colocalization analysis, C57B6/J mice were injected with choleratoxin B (CTB) conjugated to Alexa-488 in the NAc (400nl) and CTB conjugated to Alexa-647 in the OFC (300nl). For 5-HT GRAB sensor 2-photon imaging, C57B6/J mice were injected with a virus expressing 5HT3.0 GRAB sensor^56^ (AAV9-hsyn-5HT3.5; WZ Biosciences). For projection specific RNA sequencing, a retrograde AAV expressing cre recombinase under the TPH2 promoter^57^ (rAAV TPH2-cre-WPRE-hGH poly A; BrainVTA PT-0396) was injected into either the NAc or OFC of RiboTag mice. For pain management, adult mice received ketoprofen (5mg/kg i.p.) for up to 3 days following surgery. Recovery period for GRAB sensor and CTB surgery was 1 week. Recovery period for circuit specific expression was 6 weeks. For P9 stereotaxic injections, mice were anesthetized by isoflurane and secured in a mouse neonatal stereotaxic adaptor (Stoelting #51625). The skull was punctured at the injection site using a custom beveled Hamilton syringe (32 gauge, beveled #87908). An AAV expressing the 5-HT 2C receptor (**Extended Data Fig. 4a;** mIBA1-mHTR2C-FLAG-WPRE-miR-9-Tx4; AAVnerGene Inc.) or GFP control (**Fig. 4a** mIBA1-eGFP-WPRE-miR-9-Tx4; AAVnerGene Inc.) was delivered to the NAc (at 10° angle: AP +0.5mm, ML ± 1.2 mm, DV −2.5 mm; 400 nl/hemisphere). These viral constructs incorporate the mouse IBA1 promoter together with miRNA target sequences to restrict expression to microglia and minimize off-target transduction^58, 59^.

### Acute slice preparation

Mice were deeply anesthetized with isoflurane before rapid decapitation and brain removal. 300 µm coronal sections were collected in ice cold modified artificial cerebral spinal fluid (aCSF: 194mM sucrose, 30mM NaCl, 4.5mM KCl, 1mM MgCl_2_, 26mM NaHCO_3_, 1.2mM NaH_2_PO_4_, and 10mM D-glucose bubbled with 95% oxygen, 5% carbon dioxide) using a Leica VT1200 vibrating microtome. Brain sections were then transferred to regular aCSF (124mM NaCl, 4.5mM KCl, 2mM CaCl_2_, 1mM MgCl_2_, 26mM NaHCO_3_, 1.2mM NaH_2_PO_4_, and 10mM D-glucose, bubbled with 95% oxygen, 5% carbon dioxide) and incubated at 32°C for 30 min for recovery, followed by room temperature incubation until recording following previously established methods^60, 61^.

### Whole cell patch clamp electrophysiology

Slices were placed into a recording chamber and perfused with temperature-controlled aCSF (29-31°C) during the recording procedure (aCSF: 124mM NaCL, 4.5mM KCl, 1mM MgCl_2_, 26mM NaHCO_3_, 1.2mM NaH_2_PO_4_, 10mM D-glucose, 2mM CaCl_2_, bubbled with 95% oxygen, 5% carbon dioxide). Virally labeled serotonergic neurons in the DRN were fluorescently visualized using Eclipse FN1 Nikon microscope and X-Cite fluorescence illumination system (series 120Q). Images were digitally rendered using a DAGE-MTI digital camera and VLC media player (Softonic). For current clamp recordings, borosilicate glass recording pipettes (3-7MΩ resistance) were filled with a potassium gluconate-based solution (126mM potassium gluconate, 4mM KCl, 10mM Hepes, 4mM ATP-Mg, 0.3mM GTP-Na and 10mM phosphocreatine). For voltage clamp recordings, dorsal raphe serotonergic neurons were voltage-clamped at −65mV using a MultiClamp 700B Amplifier (Molecular Devices). Miniature excitatory postsynaptic currents (mEPSCs) were pharmacologically isolated by adding tetrodotoxin (1µM) and picrotoxin (50µM) to the recording aCSF. Miniature EPSCs were recorded using a borosilicate glass pipette (3-7MΩ resistance) filled with cesium-based internal solution (120mM CsMeSO_3_, 10mM HEPES, 5mM NaCl, 10mM TEA-Cl, 1.1mM EGTA, 0.3mM Na-GTP, 5mM QX-314, 4mM Mg-ATP) and 2-4% biocytin. Miniature inhibitory post synaptic currents (mIPSCs) were pharmacologically isolated by adding tetrodotoxin (1µM), DL-AP5 (50µM) and CNQX (5µM) to the recording aCSF. Miniature IPSCs were recording using a borosilicate glass pipette (3-5 MΩ resistance) filled with a high chloride-based internal solution (150mM CsCl, 10mM HEPES, 2mM MgCl_2_, 0.3mM Na-GTP, 5mM QX-314, 3mM Mg-ATP, and 0.2mM BAPTA) and 2-4% biocytin. Signals were filtered at 2 kHz, digitized at 10 kHz and acquired using Clampe× 10.4.1.4 software (Molecular Devices). Data were analyzed offline using Mini Analysis Program (Synaptosoft). Sections with biocytin filled neurons were post-fixed in paraformaldehyde (4%) for 24h.

### Sholl analysis

Image processing and analysis were conducted using previously established methods^62^. Briefly, confocal images of dye-filled neurons were put through an Ilastic pixel segmentation pipleline to generate binary segmentation images^63^. Binary images were processed using custom python scripts based on the sholl analysis FIJI plugin that skeletonizes the image and plots concentric rings (30um step size) from the soma of each cell. All code for 2D Sholl analysis is available on GitHub: https://github.com/bendevlin18/sholl-analysis-python.

### 3D projection-specific circuit mapping

Immunofluorescent images were converted to PNG images using ImageJ and subsequently preprocessed, registered and analyzed using QUINT workflow^64^. The spatial registration of the mouse brains was performed using QuickNII^65^ with Allen Mouse Brain Atlas version 3 2017 reference atlas. Subsequently, images were processed with VisuAlign and registered in QuickNII. Ilastik software was used for pixel classification of the images. MeshView Atlas Viewer was used to visualize the results in 3D atlas space.

### RNA sequencing

RNA was isolated from RiboTag mice using previously established methods^66^. Briefly, DRN samples were dissected and flash frozen on dry ice. Tissue was homogenized in ice-cold homogenization buffer supplemented with cycloheximide and RNase inhibitors using Dounce homogenizer. Homogenate incubated with anti-HA antibody for 4h at 4C. Magnetic Dynabeads were added to the lysate-antibody mixture and incubated at 4C for 2h. Ribosomes were isolated using magnetic rack and RNA was isolated using TRIzol RNA extraction reagent. RNA samples were sent to Plasmidsaurus for 3’ tag Illumina sequencing and data analysis following their standard bioinformatics pipeline. FastQ files were generated with BCL Convert (v4.3.6). Read-filtering using FastP (v0.24.0) using a minimum Phred quality score of 15 and a minimum length requirement of 50bp. Alignment to the mouse genome using STAR aligner (v2.7) with non-canonical splice junction removal and output of unmapped reads. UMI based de-duplication to remove PCR and optical duplicates using UMICollapse (v1.1.0). Alignment quality metrics, strand specificity, and read distribution across genomic features using RustQC (v0.2.1). Gene-expression quantification using featureCounts (Subread package v2.2.2) with strand-specific counting. Differential expression was generated using edgePython (v0.2.5) and functional enrichment analysis using gene set enrichment analysis GSEApy (v0.12) using the MSigDB Hallmark gene set. Samples were excluded based on minimum of 3 million reads^67^.

### 2-photon imaging

Slices were placed into a recording chamber and perfused with temperature-controlled aCSF (29-31°C) during the recording procedure (aCSF: 124 mM NaCL, 4.5 mM KCl, 1 mM MgCl_2_, 26 mM NaHCO_3_, 1.2 mM NaH_2_PO_4_, 10 mM D-glucose, 2 mM CaCl_2_, bubbled with 95% oxygen, 5% carbon dioxide). Images were collected from a Bruker ultima 2P plus with an Olympus Mplan 20x immersion objective using Prarie view software (version 5.6). InSight Tunable laser (Spectraphysics) was used to excited viral fluorophores (950nm). For 5-HT GRAB sensor recordings, images were acquired at 8.5Hz on 128×128 frame with a 3x digital zoom. A Grass SD9 stimulator was used to deliver 3 sec pulses of depolarizing current (4-20Hz) through a bipolar concentric electrode (FHC #30200) placed at the surface of the tissue near the anterior commissure of the nucleus accumbens. For microglia GCaMP recordings, tetrodotoxin (1 µM) was added to the bath solution to inhibit neuronal activity. 20 µM z-stack images (1 µM steps) were collected every minute. Following a 10-minute baseline recording, the selective 5-HT 2C receptor agonist (WAY-161503; 10 µM) was washed onto the slice for 15 minutes.

### Microglia isolation

Mice were anesthetized with CO_2_ and transcardially perfused with saline to remove circulating monocytes. Nucleus accumbens (NAc) and orbitofrontal cortex (OFC) were microdissected and microglia were subsequently isolated using CD11b-beads according to our previously published protocol^68^

### qRT-PCR

RNA was isolated using TRIzol RNA extraction reagent chloroform extraction. Isolated microglia RNA was reverse-transcribed into cDNA using a Qiagen QuantiTect Reverse Transcription kit (205311). qPCR was performed on a QuantStudio 3 thermocycler (Applied biosystems) using Sybr/Rox amplification with a QuantiFAST PCR kit (204056). Primer sequences are listed in table 5. Fold change was calculated using the 2^(−ΔΔCT)^ method with 18S as an endogenous control.

### Operant reversal learning

The lever acquisition and reversal learning task were adapted from previous methods^69^. Mice were single housed and food restricted to 85-90% baseline body weight to motivate operant responding. Mie were first familiarized with ∼10 reward pellets/mouse in the home cage. The following day, mice were placed in sound attenuated operant boxes equipped with two retractable levers and a food pellet dispenser (MED-307A-D1; MedAssociates). During initial lever acquisition, mice learned to lever press on an FR1 schedule for 14 mg dustless pellets (#F05684, BioServ, Frenchtown, NJ). For each training session, mice were given 50 lever presses or 30mins before the session ended. After two consecutive days of >40/50 lever presses (>80%), mice transitioned to the single discrimination phase. During this phase, the house lights of the operant box were turned off, and a probe light illuminated above a single correct lever (left or right) determined by the animals natural side preference during initial lever acquisition. The criterion to pass the single discrimination phase was two consecutive days of >80% performance accuracy. Afterward, mice transitioned into the reversal learning phase, where pressing the previously rewarded lever resulted in a time out (10 second period, no pellet reward) and the previously unrewarded lever resulted in the delivery of a pellet reward. Mice performed the reversal task for 5 consecutive days until performance stabilized near 80% correct lever presses demonstrating a clear update in the reward contingency.

### Voluntary ethanol consumption

Mice were group housed in a reverse light cycle room (lights off at 1000h, on at 2200h). The drinking in the dark (DID) protocol was adapted from Rhodes et al.^70^ and our previously established methods^71^. Briefly, mice were given two hours of access to the ethanol drinking bottle in the first three days, and on day four, mice were given four hours of access. Bottle weights were recorded to determine total consumption and adjusted to body weight (g/kg). For the two-bottle choice assay, mice were given a two-bottle choice test of water and ethanol (20%) for a 2-h duration to determine drinking preference. Bottle weights were recorded to determine total consumption and adjusted to body weight (g/kg). Mice received an injection of deschloroclozapine^72^ (0.1mg/kg i.p.) 45min prior to the two-bottle choice test.

### Circadian behavior

Mice were individually housed in cages (35.3 x 23.5 x 20 cm) equipped with 12.7 cm diameter running wheel (Lafayette Instrument #80820FS). Water and rodent chow (Lab Diet 5001; Lab Diet) were provided throughout the course of the study. Activity data were collected using the Scurry Activity Monitoring software (Lafayette Instrument #86165). Mice were placed on a 12:12 light-dark cycle (lights off at 0800h, on at 2000h) for 10 days. Following the entrainment to the 12:12 cycle, mice were placed into constant darkness (free run) for 24 days. On day 24, mice were subjected to a 6h light pulse at CT 12 based on the individual “tau” for each animal and returned to constant darkness for another 13 days. On day 38, mice were placed back on a 12:12 light-dark cycle for 8 days for a final entrainment. Actograms were generated in excel.

### Conditioned place preference test

The following was adapted from Morales and colleagues^44^. Briefly, mice were food-restricted to 85-90% of their baseline body weight. Mice were tested in a two-chamber CPP box (MED Associates, Inc.) and time spent in each chamber was recorded using Med Associates MED-PC software. On day 1, mice were allowed to freely explore both chambers for 20 minutes to establish a baseline chamber preference. On day 2 mice were confined to the preferred chamber for 20 minutes with no food. On day 3 mice were confined to their non-preferred chamber for 20 minutes with a food pellet. On day 4, mice were allowed to move freely between both chambers for 20 minutes. For the DREADDs study, on day 2 mice received a vehicle injection (0.5% DMSO in saline, i.p.) 45 minutes prior to conditioning. On day 3, mice received an injection of deschloroclozapine^72^ (0.1mg/kg i.p.) 45min prior to conditioning.

### Statistical analysis

All statistical analyses were performed in GraphPad Prism 8.0. Data are represented as mean ± SEM. Sample size and statistical analysis are defined in the figure legend. For image analysis and electrophysiology data, sample size is defined by an individual animal consisting of an average of multiple images or cells per animal and is represented graphically by gray dots. All diet offspring data were collected across a minimum of 3 litters to account for potential litter effects. Comparisons between two groups were analyzed with a Student’s *t* test while comparisons of three or more groups were analyzed with two-way ANOVA. Multiple comparisons were corrected with Sidak’s test or Dunnett’s test when comparing multiple time points to a single baseline.

## Results

We examined the developmental trajectory of serotonin fibers in two anatomically and developmentally distinct brain regions: the orbitofrontal cortex (OFC) and the nucleus accumbens (NAc) (**Fig. 1a-d**). These regions were selected to capture cortical and subcortical targets of anatomically heterogenous dorsal raphe nucleus (DRN) projections^73, 74^. We first quantified serotonergic innervation across the first three postnatal weeks in mice on standard chow, a period marked by robust axonal expansion and refinement corresponding with maturation of target brain regions^53, 75^. Serotonin fibers were visualized using an antibody against the serotonin transporter (SERT), and fiber density was quantified as percent area coverage^76^. In the NAc, serotonergic innervation peaked at postnatal day (P)14 (**Fig 1c**). In contrast, serotonergic innervation of the OFC followed a delayed trajectory, with maximal fiber density observed at P21 (**Fig. 1d**), consistent with protracted development of cortical monoaminergic circuits^77–79^.

**Figure 1.**
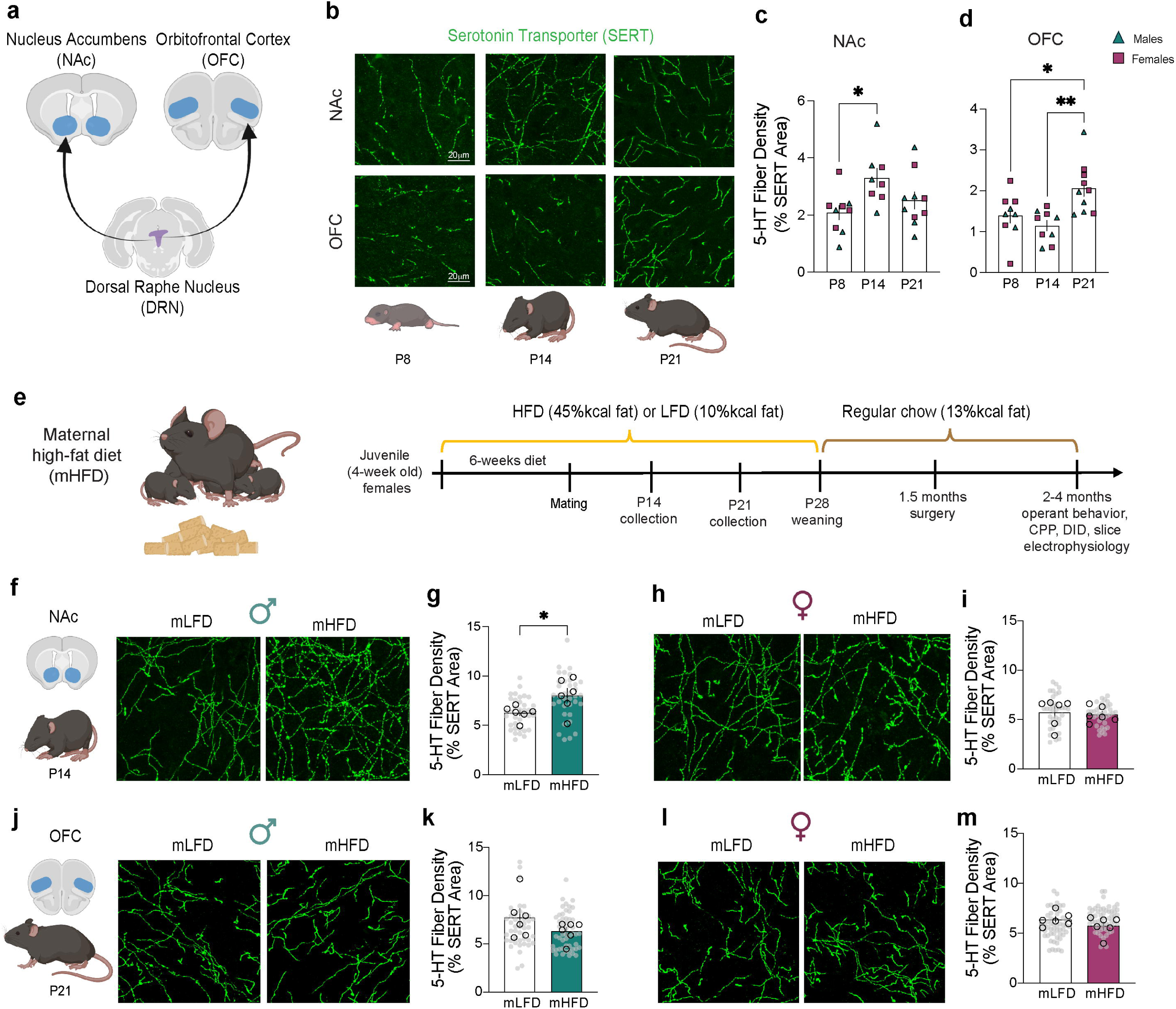
Maternal high-fat diet selectively alters serotonergic innervation of the nucleus accumbens in male offspring. **a,** Schematic illustrating serotonergic projections from the dorsal raphe nucleus (DRN) to the nucleus accumbens (NAc) and the orbitofrontal cortex (OFC). **b,** Representative confocal images of serotonergic fiber innervation in the NAc and OFC across postnatal development (P8-P21). **c,** Serotonergic fiber innervation in the NAc peaks at P14 in both male and female mice (n=4 males, 4 females; p<0.05; One-way ANOVA, Tukey’s multiple comparison test). **d,** In the OFC, serotonergic fiber innervation peaks at P21 (n=4 males, 4 females; p<0.01; One-way ANOVA, Tukey’s multiple comparison test). **e,** Schematic and experimental timeline of maternal high-fat diet (mHFD) exposure. **f-g**, mHFD increases serotonergic innervation to the NAc at P14 in male offspring (n=6 mLFD, 6 mHFD; p<0.05; Student’s t-test), **h-i** but not in female offspring (n=6 mLFD, 6 mHFD offspring; p=0.74; Student’s t-test). **j-m**, mHFD does not alter serotonergic innervation to the OFC at P21 in male (n=6 mLFD, 6 mHFD; p=0.18; Student’s t-test) or female offspring (n=6 mLFD, 6 mHFD; p=0.29; Student’s t-test). Data represented as mean ± SEM. Gray points denote individual images. *p<0.05.

We next assessed the impact of maternal high-fat diet (mHFD) on serotonergic innervation to these two brain regions (**Fig. 1e**). Consistent with our previous findings^11^, mHFD increased offspring body weight at P14 in both males and females compared to maternal low-fat diet (mLFD) offspring (**Extended data Fig. 1a**). This effect persisted through P21 (**Extended data Fig. 1b**). mHFD produced a sex-and region-specific effect on serotonergic innervation. Specifically, male mHFD offspring exhibited increased serotonergic fiber density in the NAc at P14 (**Fig. 1f,g**). This effect was not observed in female offspring (**Fig. 1h,i**). By contrast, mHFD did not alter serotonergic innervation of the OFC at P21 (**Fig. 1j-m**) or P14 (**Extended data Fig. 1c-f**) in either males or female offspring. When we measured serotonergic fiber innervation to the NAc at P21, we no longer observed hyperinnervation in mHFD male offspring (**Extended data Fig. 1g,h**) and no effect was observed in females at this age (**Extended data Fig. 1i,j**). Overall, we identified a male-specific impact of mHFD on serotonergic fiber density in the NAc at P14.

Provided the essential role of microglia in sculpting developing neural circuits^31, 32, 34–38^, we next assessed whether mHFD impacts microglial phagocytosis of serotonergic fibers in the NAc and OFC, focusing on male offspring where we observed serotonergic hyperinnervation (**Fig. 2**). At P14, NAc microglia of mHFD male offspring exhibited reduced lysosomal volume, measured by CD68 immunoreactivity, compared with mLFD microglia (**Fig. 2a,b**). Consistent with decreased lysosomal volume, NAc microglia from mHFD male offspring engulfed less SERT-positive material than those from mLFD males (**Fig. 2c**), paralleling the serotonergic hyperinnervation observed in the NAc at this age. By P21, lysosomal volume in NAc microglia from mHFD males was increased relative to mLFD (**Fig. 2d,e**). However, SERT phagocytosis remained reduced (**Fig. 2f**), indicating a persistent deficit in serotonergic fiber engulfment across this developmental window. In female offspring, mHFD produced a trend toward reduced lysosomal volume in NAc microglia at P14 (**Extended data Fig. 2a,b**) and did not alter SERT phagocytosis (**Extended data Fig. 2c**). No effect of mHFD on NAc microglial lysosomal volume or SERT phagocytosis was observed in females at P21(**Extended data Fig. 2d-f**).

**Figure 2.**
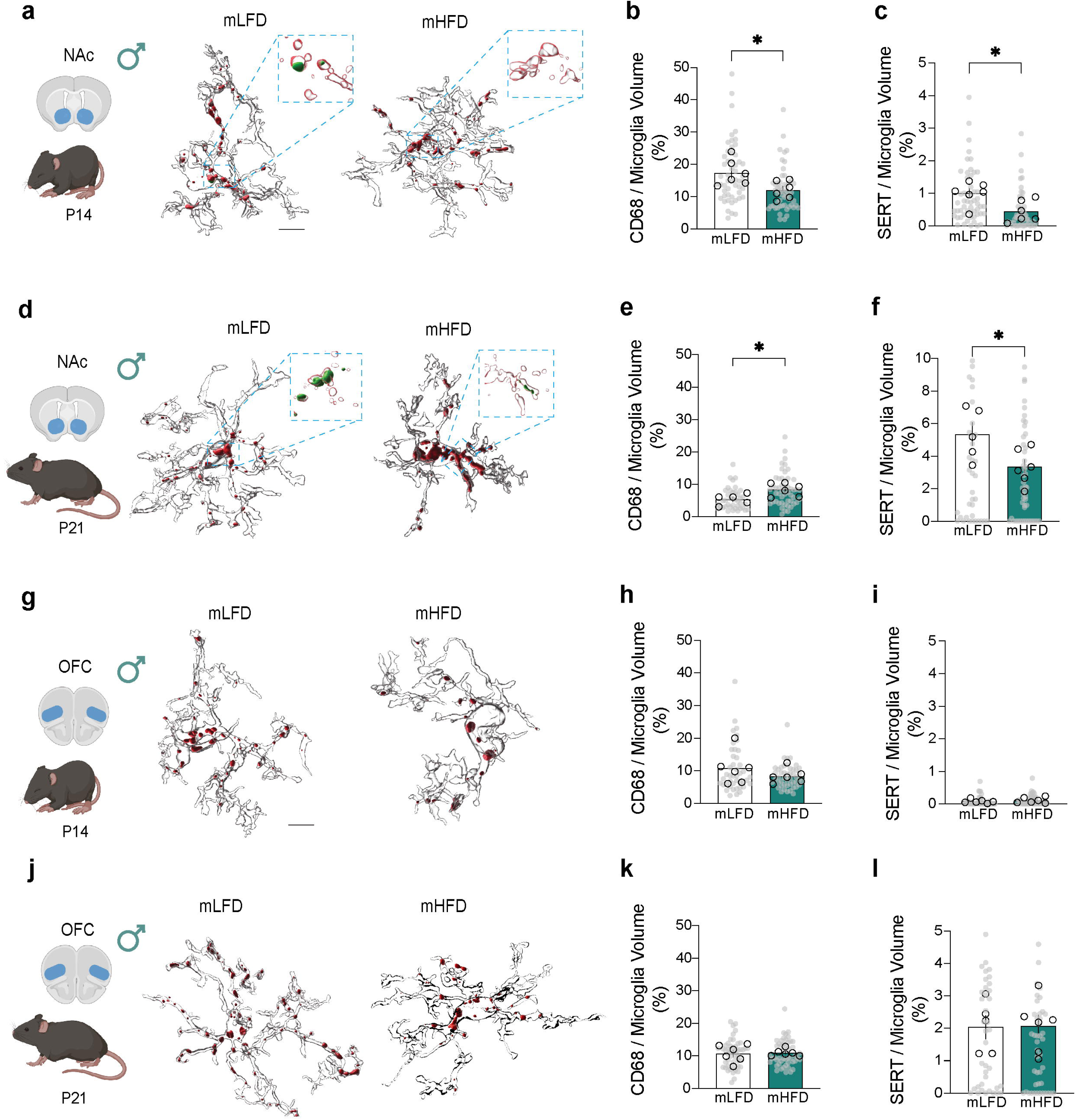
Maternal high-fat diet reduces microglial phagocytosis of serotonergic fibers in the male nucleus accumbens. **a**, Representative 3D reconstructions of NAc microglia from mLFD and mHFD male offspring at P14, showing lysosomal content (CD68, red) containing serotonergic material (SERT, green). Scale bar 8µm. **b**, NAc microglia from mHFD male offspring exhibit reduced lysosomal volume compared with mLFD microglia at P14 (n=6 mLFD, 6mHFD offspring; p<0.05; Student’s t-test). **c**, NAc microglia from mHFD males engulf less SERT-positive material than those from mLFD males at P14 (n=6 mLFD, 6mHFD offspring; p<0.05; Student’s t-test). **d**, Representative 3D reconstructions of NAc microglia from mLFD and mHFD male offspring at P21. **e**, NAc microglia from mHFD male offspring exhibit increased lysosomal volume relative to mLFD microglia at P21(n=6 mLFD, 6mHFD offspring; p<0.05; Student’s t-test). **f**, Despite increased lysosomal volume, SERT phagocytosis remains reduced in NAc microglia from mHFD males at P21 (n=6 mLFD, 6mHFD offspring; p<0.05; Student’s t-test). **g**, Representative 3D reconstructions of OFC microglia from mLFD and mHFD male offspring at P14, showing CD68-positive lysosomes (red) and SERT (green). Scale bar 8µm. **h**,**i**, mHFD does not alter lysosomal volume (**h**, n=6 mLFD, 6mHFD offspring; p=0.32; Student’s t-test), or SERT phagocytosis (**i**, n=6 mLFD, 6mHFD offspring; p=0.49; Student’s t-test) in OFC microglia from P14 male offspring. **j**, Representative 3D reconstructions of OFC microglia from mLFD and mHFD male offspring at P21. **k,l,** No effect of mHFD on lysosomal volume (**k**, p=0.86) or SERT phagocytosis (**l**, p=0.94) in OFC microglia from P21 male offspring (n=6 mLFD, 6mHFD offspring; Student’s t-test). Data represented as mean ± SEM. Gray points denote individual cells. *p<0.05.

In the OFC, mHFD did not impact lysosomal volume or SERT phagocytosis at P14 (**Fig. 2g-i**) or P21 (**Fig. 2j-l**). In female offspring, mHFD similarly had no effect on OFC microglia lysosomal volume or SERT phagocytosis at P14 (**Extended data Fig. 2g-i**). At P21, however, OFC microglia from mHFD females exhibited a significant increase in lysosomal volume compared to mLFD microglia (**Extended data Fig. 2j,k**), although SERT phagocytosis remained unchanged (**Extended data Fig. 2l**).

Microglia express serotonin receptors and serotonergic signaling modulates core microglia functions, including phagocytosis, process motility, and exosome secretion^80–84^. We therefore investigated whether microglia in the NAc and OFC exhibit distinct serotonin receptor expression that could mediate the differential outcomes we observed in mHFD offspring. Using our established microglial isolation method^68^ (**Fig. 3a**), we found that at P14, microglia from the NAc of both male and female offspring expressed higher levels of *Htr2c* and lower levels of *Htr1b* compared with OFC microglia (**Fig. 3b**). No regional differences were observed in expression of *Htr1a*, *Htr2a*, or *Htr2b* expression in microglia at this age (**Fig. 3b**). In non-microglial (CD11B negative) cells, we similarly observed region-specific differences in serotonin receptor expression at P14. Non-microglial cells from the NAc exhibited higher expression of *Htr2c*, *Htr1b* and *Htr2b*, whereas OFC cells showed higher expression of *Htr1a* and *Htr2a* (**Extended Data Fig. 3a**). Some, but not all, of these regional differences were still evident in adulthood (2-4 months). Adult NAc microglia continued to exhibit higher *Htr2c* expression in both sexes, with no differences in *Htr2b* expression between regions (**Extended Data Fig. 3b**). In contrast, adult OFC microglia displayed higher expression of *Htr1a* and *Htr2a*, whereas regional differences in *Htr1b* expression were no longer evident (**Extended Data Fig. 5b**). Non-microglial cells maintained a similar pattern of regional receptor expression across development, with higher *Htr1a* and *Htr2a* expression in the OFC and higher *Htr1b*, *Htr2b*, and *Htr2c* expression in the NAc (**Extended Data Fig. 3c**).

**Figure 3.**
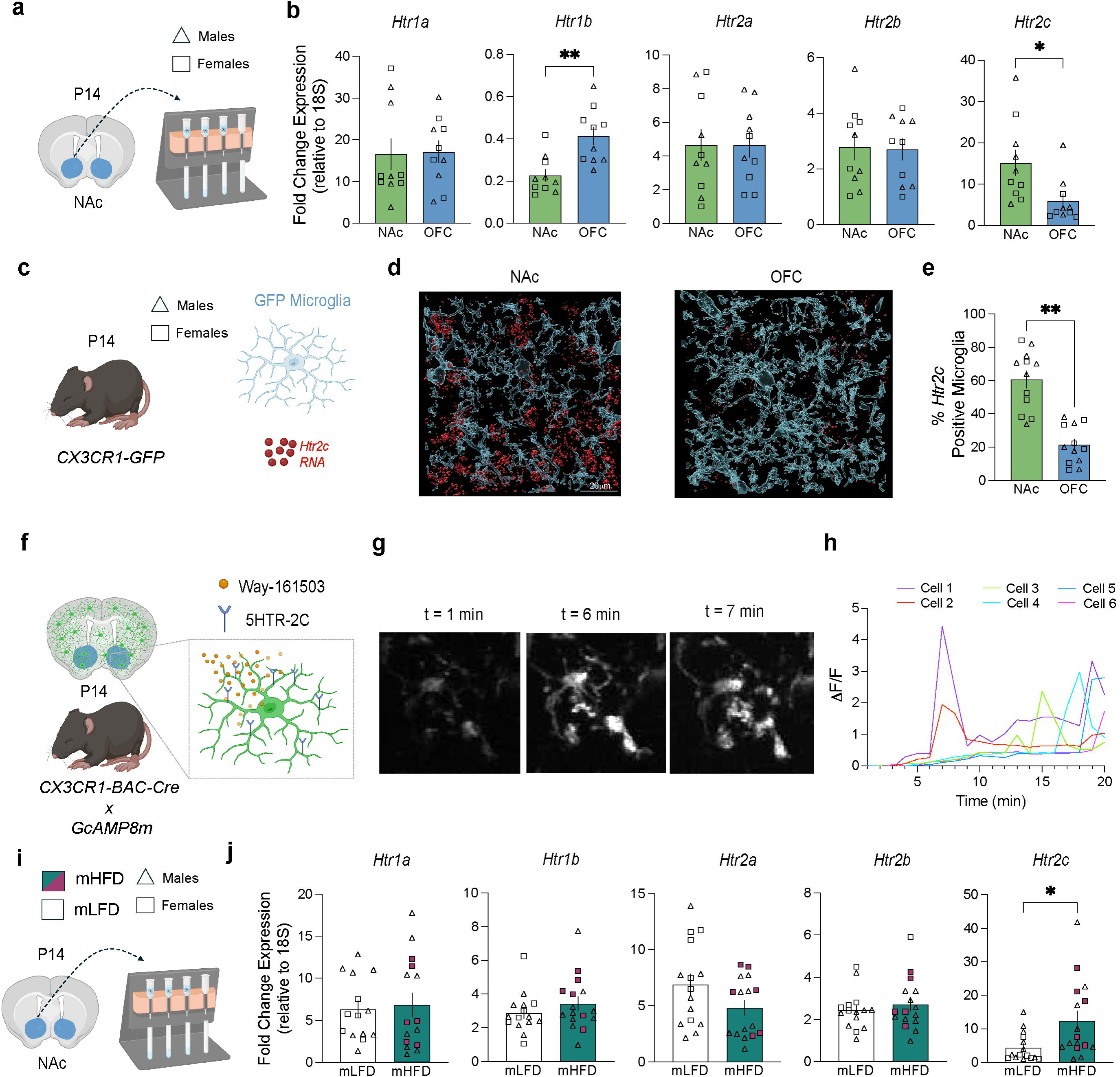
Regional heterogeneity of microglial serotonin receptor expression in the orbitofrontal cortex and nucleus accumbens. **a**, Schematic illustrating microdissection of the NAc and OFC followed by microglia isolation using magnetic cell separation. **b**, Microglia isolated from the NAc express higher levels of *Htr2c* (p<0.05) and lower levels of *Htr1b* (p<0.01) compared with OFC microglia (n=5 males, 5 females per brain region; Student’s t-test). No regional differences were observed in expression of *Htr1a* (p=0.9), *Htr2a* (p=0.74), or *Htr2b* (p=0.88). **c**, Schematic depicting the CX3CR1-GFP mouse line used to label microglia and RNA fluorescence in situ hybridization (RNA-FISH) probes targeting *Htr2c*. **d**, Representative 3D reconstructions of microglia (blue) with *Htr2c* RNA puncta in the NAc (left) and OFC (right). **e**, A greater proportion of microglia in the NAc express *Htr2c* RNA puncta compared with OFC microglia (n=6 males, 6 females per brain region; p<0.01, Student’s t-test). **f**, Schematic illustrating *CX3CR1-BAC-Cre* crossed with *GcAMP8m-floxed* mice to monitor microglial calcium dynamics in response to bath application of the selective 5HTR-2C agonist Way-161503 (10µM). **g**, Representative two-photon time-lapse images of NAc microglia at baseline (1min) and following application of Way-161503 (6-7min). **h**, NAc microglia exhibit increased calcium activity following Way-161503 application relative to baseline (n=3 males, 3 females; p<0.01, paired t-test). **i**, Schematic outlining mHFD exposure and P14 microglia isolation for serotonin receptor expression analysis. **j**, mHFD selectively alters microglial *Htr2c* receptor expression in both males and female P14 offspring (n=9 males, 6 females per diet; p<0.05, Student’s t-test). No effects of mHFD were observed on expression of *Htr1a* (p=0.76), *Htr1b* (p=0.29) or *Htr2b* (p=0.49), and a non-significant trend toward reduced *Htr2a* expression was observed (p=0.09). Data represented as mean ± SEM. *p<0.05, **p<0.01.

We focused subsequent analysis on the 5HTR-2C receptor, given the persistent enrichment in the NAc across development and recent work demonstrating activation of a class II serotonin receptor, 5HTR-2B, regulates microglial phagocytosis in the hippocampus^82^. We first confirmed elevated *Htr2c* expression in NAc microglia using RNA fluorescence in situ hybridization (RNA-FISH) in CX3CR1-GFP mice (**Fig. 3c,d**). Approximately 60% of NAc microglia express *Htr2c*, compared to 20% of OFC microglia (**Fig. 3e**). Because 5HTR-2C is Gq coupled, its activation engages PLC-IP3 signaling to elevate intracellular calcium. We therefore tested whether microglial 5HTR-2C are functionally active by monitoring calcium dynamics using GcAMP8M-floxed mice crossed to CX3CR1-BAC-cre mice (**Fig. 3f**). Using 2-photon imaging, we recorded baseline microglial calcium activity in the NAc and then applied the selective 5HTR-2C agonist Way-161503 (10µM). Bath application of WAY-161503 elicited a robust increase in microglial GCaMP fluorescence relative to baseline (**Fig. 3g,h**).

We next asked whether mHFD alters serotonin receptor expression in NAc microglia (**Fig. 3i**). We found that mHFD selectively increased *Htr2c* expression in NAc microglia of P14 offspring, with no detectable effects on expression of *Htr1a*, *Htr1b*, or *Htr2b* in either male or female offspring (**Fig. 3j**). Similarly, we observed no impact of mHFD on expression of *Htr1a*, *Htr1b*, *Htr2a*, or *Htr2b* in non-microglia (CD11B negative) cells in the NAc (**Extended Data Fig. 3d**).

To determine if 5HTR-2C expression impacts microglia phagocytosis, we developed a novel viral strategy to genetically modify microglia. We used a viral plasmid containing the mouse IBA1 promoter^59^ and micro RNAs and introduced loxP sites to further reduce off-target expression in neurons (**Fig. 4a**). We crossed our Cx3CR1-BAC-cre to SELECTIV mice (**Fig. 4b**), which over expresses the AAV receptor^85^ to facilitate viral transduction in notoriously difficult cells, such as microglia^86, 87^. Using this approach, we observed robust microglia transduction of AAV-GFP in P14 offspring (**Fig. 4c-e**) that was highly restricted to IBA1 positive microglia (**Extended Data Fig. 4a,b**). We next transduced NAc microglia with an AAV plasmid expressing the 5HTR-2C (**Extended Data Fig. 4c**) to phenocopy the increased expression observed in mHFD offspring (**Fig. 3j**). Using RNA-FISH, we observed greater *Htr2c* puncta volume in IBA1-positive microglia in the NAc in both male and female offspring (**Extended Data Fig. 4d,e**). AAV transduction of 5-HTR2C in microglia was sufficient to increase serotonergic innervation in the NAc compared to GFP control treated offspring (**Fig. 4f,g**).

**Figure 4.**
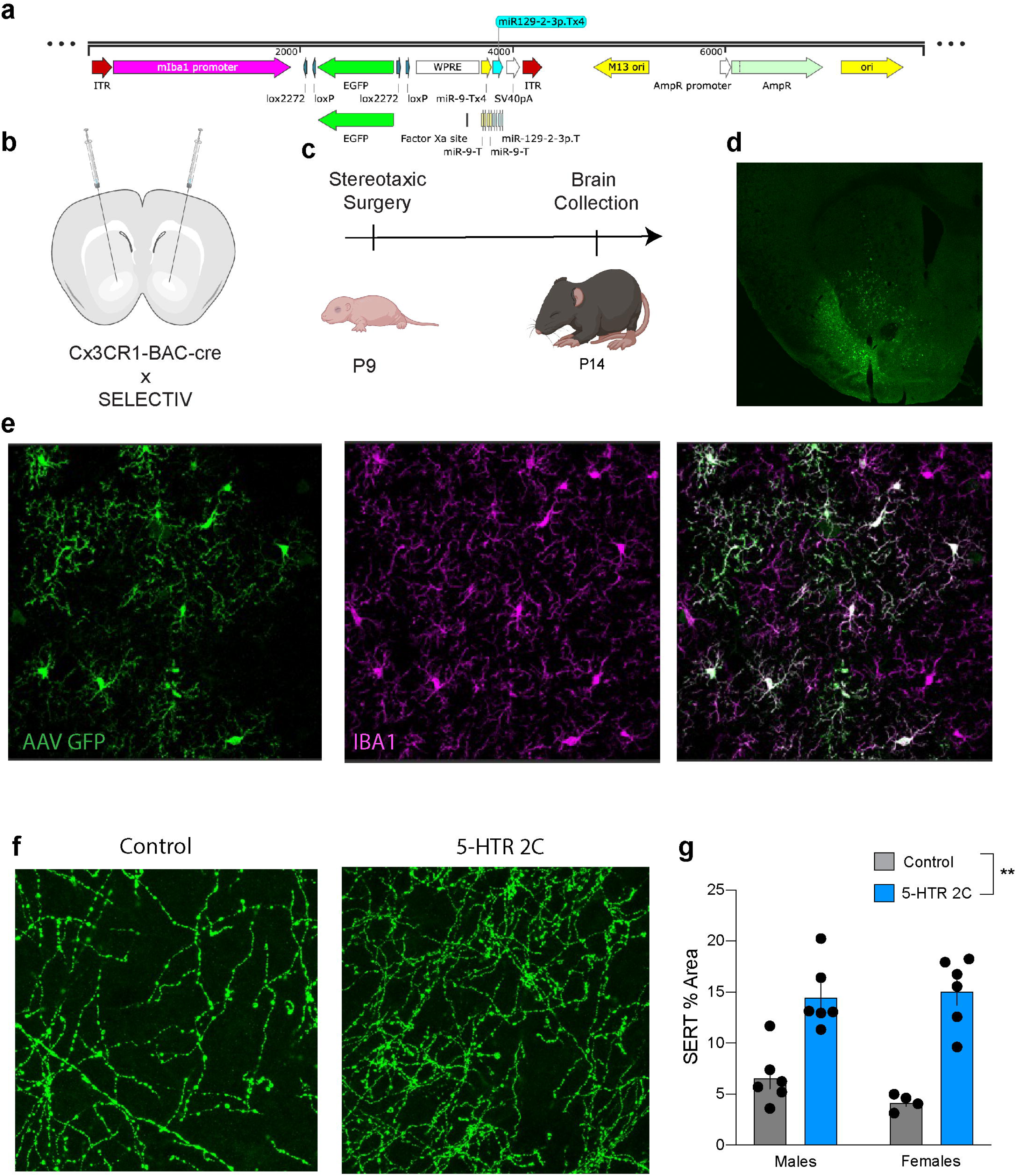
Microglia overexpression of the 5-HT2C receptor increases serotonin fiber innervation in the nucleus accumbens. **a**, Schematic of microglia-targeting AAV plasmid expressing GFP under the mouse IBA1 promoter. **b**, Schematic of stereotaxic surgery in Cx3CR1-BAC-cre:SELECTIV mice to genetically manipulate microglia in the Nac. **c**, Timeline depicting stereotaxic surgery in P9 pups and brain collection at P14. **d**, Representative 10x confocal image depicting P14 viral expression of GFP in NAc microglia. **e**, Representative 30x confocal images demonstrating AAV GFP expression in IBA1 positive microglia in the NAc. **f**, Representative confocal images of serotonin fibers (SERT) from P14 offspring treated with AAV GFP control or 5-HT2C overexpression. **g**, Both male and female offspring treated with 5-HT2C expressing AAV exhibited increased serotonin innervation in the NAc (n=5 male + 4 female controls, 6 male +6 female 5-HT2C; main effect of treatment p< 0.01; two-way ANOVA). Data represented as mean ± SEM. **p<0.01.

The basis for the selective impact of mHFD on DRN→NAc, but not DRN→OFC, serotonin neurons may also reflect heterogeneity among DRN serotonin neuron subtypes^73, 88, 89^. Previous studies have described spatial segregation of serotonin neurons along the dorsal-ventral axis of the DRN, with dorsal neurons preferentially targeting subcortical forebrain regions and ventral neurons projecting to cortical areas^73, 74^. To test whether such spatial organization underlies the projection-specific effects observed here, we mapped the distribution of NAc and OFC projecting serotonin neurons within the DRN using an intersectional viral strategy (**Extended Data Fig. 5a,b**). Confocal images of coronal DRN sections were registered to a 3D Allen Brain Atlas and contrary to expectations, we observed no dorsal-ventral segregation of the NAc versus OFC projecting serotonin neurons (**Fig. 5a,b**). We next asked whether NAc and OFC projecting serotonin neurons represent distinct cellular populations. To this end, we used the retrograde neuronal tracer cholera toxin subunit B (CTB) to fluorescently label OFC and NAc projecting neurons in the DRN (**Fig. 5 c,d**). We found that in the DRN (−4.5mm to −4.8mm from bregma), the majority of neurons were projection-target specific, with minimal collateralization to the other brain region (**Fig. 5e**).

**Figure 5.**
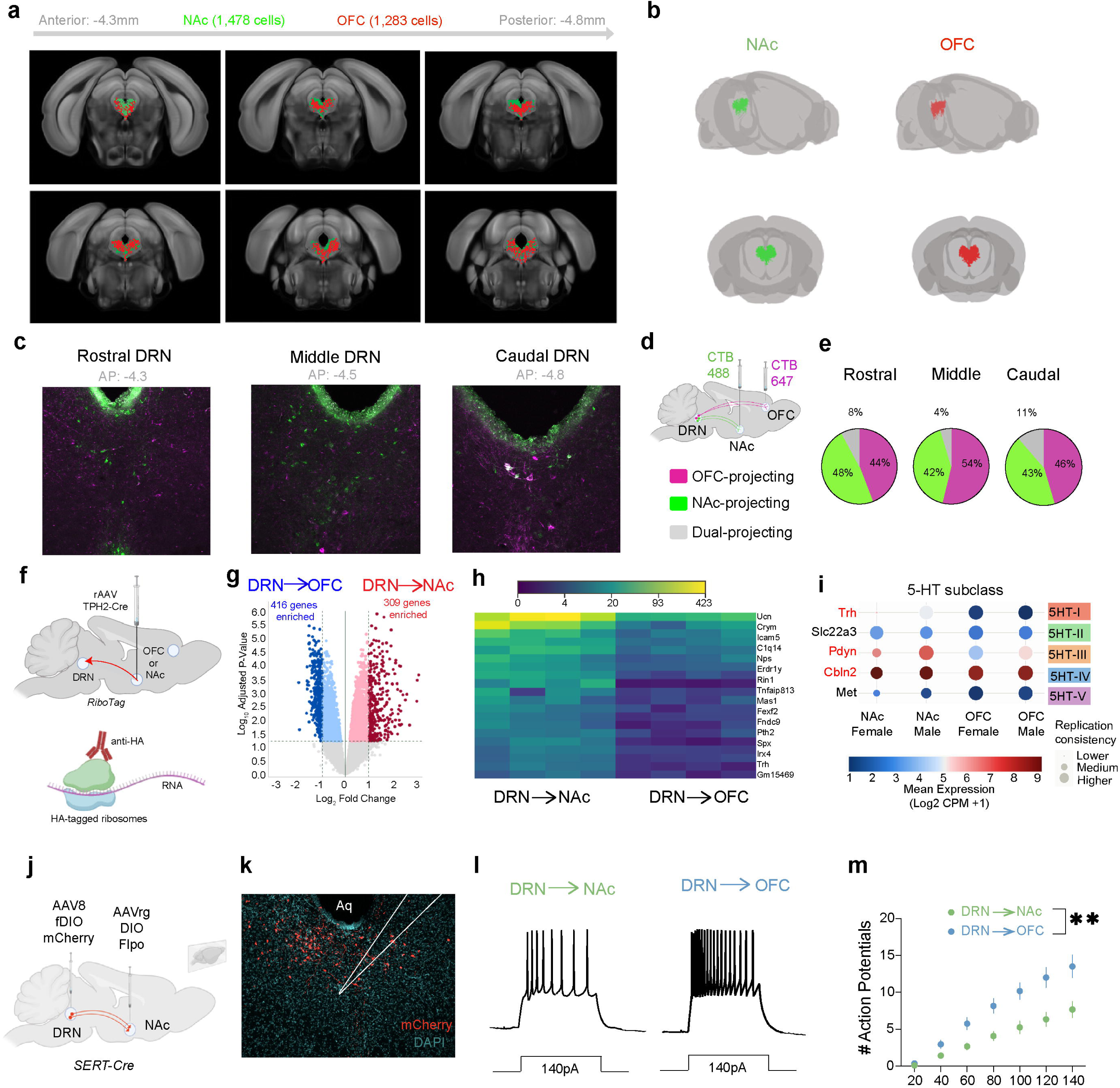
DRN→NAc and DRN→OFC serotonergic neurons are transcriptionally and functionally distinct subpopulations. **a,** Atlas-based reconstruction of the DRN (AP −4.5 to - 4.95mm) showing serotonergic neurons projecting to NAc (green) or OFC (red), labeled using an intersectional viral strategy (NAC: n=7 males, 6 females; OFC: n=6males, 6 females). **b**, 3D rendering of DRN→NAc and DRN→OFC serotonergic neurons.**c**, Representative images of rostral (AP −4.3mm), middle (AP −4.5mm), and caudal (AP −4.8mm) DRN showing OFC-projecting (magenta) and NAc-projecting (green) neurons labeled with cholera toxin B (CTB).Scale bar = 100µm. **d**, Schematic illustrating stereotaxic injection of CTB-488 into the NAc and CTB-647 into the OFC. **e**, Quantification of OFC-projecting (magenta), NAc-projecting (green) and dual-projecting neurons across the DRN. OFC and NAc projecting neurons are primarily non-overlapping across the DRN (−4.3 to −4.8mm). **f**, schematic of stereotaxic injection of retrograde AAV expressing cre-recombinase under the THP2 promoter in RiboTag mice to isolate ribosomal RNA from projection-defined DRN serotonergic neurons. **g**, volcano plot from DRN→NAc and DRN→OFC serotonergic neurons depicting 309 NAc upregulated genes and 416 OFC upregulated genes. **h**, heat plot depicting the top 20 upregulated genes in the DRN→NAc compared to DRN→OFC serotonergic neurons. **i**, dot plot depicting genes representing the five serotonin subclasses previously identified using single cell RNA sequencing. DRN→NAc neurons align with class I, III, and IV serotonin neurons. **j**, Schematic of intersectional viral strategy to label projection-defined serotonergic neurons in the DRN with AAV mCherry. **k**, Representative confocal image depicting OFC-projecting serotonin neurons in the DRN with patch-clamp pipette. **l**, Representative current-clamp traces showing action potential firing in response to depolarizing current injection (140pA) in DRN→NAc and DRN→OFC serotonergic neurons. **m**, DRN→OFC serotonergic neurons fire more action potentials in response to depolarizing current injection than DRN→NAc neurons (n=7 males, 5 females per group; p <0.01; two-way repeated measures ANOVA).

We next sequenced each projection-defined subpopulations using RiboTag mice and a retrograde AAV expressing cre-recombinase under the tryptophan hydroxylase 2 (TPH2) promoter^57^ (**Fig. 5f; Extended Data Fig. 5c,d**). RNA isolated from HA-tagged ribosomes was sequenced, and differential gene expression (DEG) analysis identified distinct transcriptional signatures between DRN→NAc and DRN→OFC serotonin neurons (**Fig. 5g,h**). We cross-referenced the identified DEGs that aligned with the five DRN serotonin subtypes previously identified with single-cell RNA sequencing^73^. DRN→NAc neurons comprised a mixture of class I, III, and IV subtypes compared to DRN→OFC neurons (**Fig. 5i**), consistent with previous findings that projection-defined neuron populations overlap with multiple molecularly defined serotonin subtypes^73^. We further observed transcriptional segregation across previously identified gene clusters, including “channels”, “Neurotransmission”, and “transcription factors” ^73^ (**Extended Data Fig. 5e-g**). We next performed whole-cell patch clamp electrophysiology on each projection-defined neuron population (**Fig. 5j,k**). DRN→OFC neurons fired more action potentials in response to depolarizing current injection compared to DRN→NAc neurons (**Fig. 5l,m**) but observed no differences in resting membrane potential or action potential threshold (**Extended Data Fig. 5h,i**). Collectively these data demonstrate both transcriptional and functional differences between these projection-defined subpopulations.

Previous studies have demonstrated that cortical-projecting DRN serotonin neurons are enriched for vesicular glutamate transporter 3 (*Vglut3*) relative to subcortical-projecting DRN serotonin neurons^73, 74, 88^. To further assess cellular heterogeneity between DRN→NAc and DRN→OFC projecting neurons, we examined expression of *Vglut3* and glutamate decarboxylase 2 (*Gad2*) using RNA-FISH. Consistent with prior reports, *Vglut3* expression was concentrated along the midline of the DRN, whereas *Gad2* expression was enriched in the lateral wings (**Extended Data Fig. 5j**). However, we detected no differences in the proportion of DRN→NAc or DRN→OFC serotonin neurons co-expressing either *Vglut3* or *Gad2* (**Extended Data Fig. 5k**).

We next examined the impact of mHFD on DRN→ NAc and DRN→OFC physiology. mHFD reduced the tonic firing rate of DRN→ NAc serotonergic neurons in male but not female adult offspring (**Fig. 6a**). In contrast, mHFD did not alter the tonic firing rate of DRN→OFC serotonergic neurons (**Fig. 6b**). mHFD also had no effect on firing evoked by depolarizing current injection in either projection-defined neuron population (**Fig. 6c,d**), nor on other measures of intrinsic excitability across sexes (**Extended Data Fig. 6a-h**). Together, these findings indicated that mHFD selectively reduces the basal activity of DRN→ NAc neurons while preserving their capacity to fire at high frequencies when this circuit is recruited (e.g., behavior).

**Figure 6.**
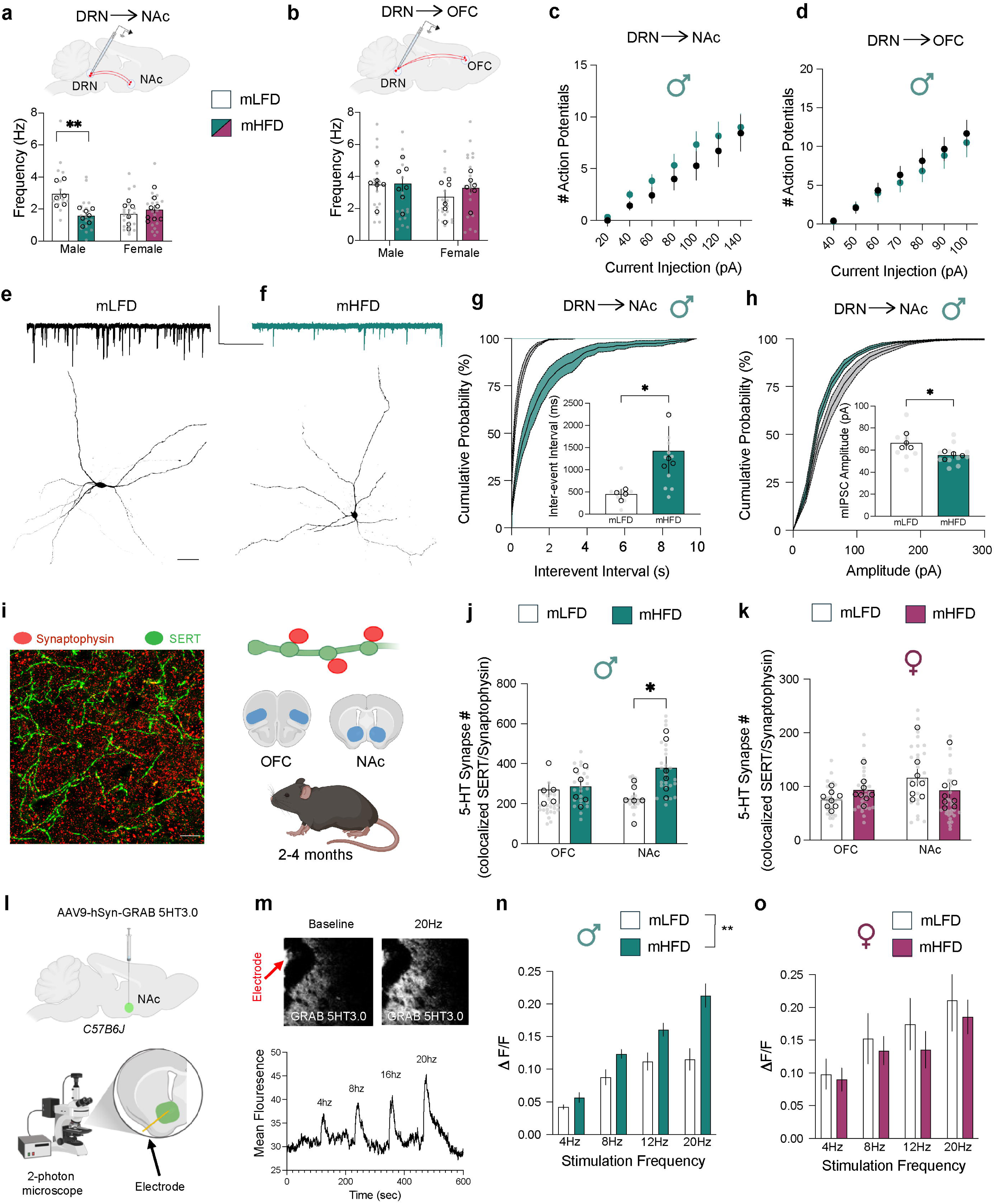
Maternal high-fat diet increases serotonin release in the nucleus accumbens of adult male offspring. **a**, mHFD reduces tonic firing rate of DRN→NAc serotonergic neurons in adult (2-4 months) male offspring (n=6 mLFD males, 6 mLFD females, 7mHFD males, 8mHFD females; Interaction p <0.01; two-way ANOVA with Sidak’s multiple comparisons correction). **b**, mHFD does not alter tonic firing rate of DRN→OFC serotonergic neurons (n= 6 mLFD males, 6 mLFD females, 7mHFD males, 6mHFD females; main effect of diet p=0.46; two-way ANOVA). mHFD does not alter firing response to depolarizing current injection in DRN→NAc (**c**; p=0.38) or DRN→OFC (**d**, p=0.37) serotonergic neurons from adult male offspring (n=6 mLFD, 6mHFD offspring per group; two-way repeated measures ANOVA). Representative dye-filled DRN→NAc serotonergic neurons from adult mLFD (**e**) and mHFD (**f**) male offspring with representative miniature inhibitory post synaptic current (mIPSC) traces above. Scale bar=200pA, 2.5s; 50µm. **g**, mHFD increases the interevent interval of mIPSCs on DRN→NAc serotonergic neurons (p<0.05; Student’s t-test). **h**, mHFD decreases the mIPSC amplitude on DRN→NAc serotonergic neurons (n= 4 mLFD, 4mHFD male offspring; p<0.05; Student’s t-test). **i**, Representative 60x confocal image of serotonergic fibers (SERT+) and the presynaptic marker synaptophysin in the NAc of adult offspring. **j**, mHFD increases serotonergic synapse density in the NAc but not the OFC, of adult male offspring (n= 11 OFC, 12 NAc; Main effect of diet p <0.05; Interaction p=0.1; Two-way ANOVA with Sidak’s multiple comparisons correction). **k**, In contrast, mHFD does not affect serotonergic synapse density in adult female offspring (n=16 OFC, 16 NAc; Main effect of diet p=0.79; two-way ANOVA). **l**, Schematic illustrating viral expression of the GRAB 5-HT sensor in the NAc and ex-vivo slice recordings of fluorescence using a two-photon microscopy. **m**, Representative two-photon images of electrically-evoked GRAB 5-HT fluorescence at baseline (left) and during 20hz stimulation (right). Below, representative fluorescence traces evoked by 4-20Hz electrical stimulation in the NAc. **n**, Adult mHFD male offspring exhibit increased serotonin release in the NAc compared with mLFD offspring (n=6 mLFD, 6 mHFD offspring; main effect of diet p <0.01; two-way repeated measures ANOVA). **o**, No effect of mHFD on serotonin release is observed in female offspring (n=6 mLFD, 6 mHFD offspring; main effect of diet p=0.6; two-way repeated measures ANOVA). Data represented as mean ± SEM. Gray points denote individual cells or images. *p<0.05, **p<0.01.

Since our previous work demonstrated that mHFD increases microglia phagocytosis of serotonin in the fetal (E14.5) DRN of male offspring, we next determined whether early-life microglial phagocytosis produced lasting changes in serotonergic neuron morphology or local synaptic transmission. To this end, we recorded miniature excitatory (mEPSC) and miniature inhibitory post synaptic currents (mIPSC) from DRN→NAc serotonin neurons and dye-filled the cells for Sholl analysis (**Fig. 6e,f**). We found no impact of mHFD on neurite complexity in DRN→NAc serotonergic neurons in male offspring (**Extended data Fig. 6i**). However, mIPSCs from DRN→NAc serotonin neurons of mHFD male offspring were less frequent and were of smaller amplitude compared to mLFD offspring (**Fig. 6g,h**). We observed a trending increase in mEPSC frequency, but no change in event amplitude in DRN→NAc serotonin neurons relative to mLFD offspring (**Extended data Fig 6j-l**). Collectively, these physiological data indicate a gain in DRN→NAc circuit activity.

We next asked whether the serotonergic hyperinnervation observed in the NAc of P14 male offspring **(Fig. 1f,g**) persisted into adulthood. Although no differences in gross serotonergic fiber density were detected by P21 in mHFD males (**Extended data Fig. 1g,h**), we reasoned that more subtle alterations at the level of serotonergic synapses might remain. To test this, we performed a refined analysis of serotonergic synapse density by quantifying colocalization of synaptophysin puncta with SERT-positive fibers^76^ (**Fig. 6i**). We found that serotonergic synapse density was significantly increased in the NAc of adult (2-4 months) male offspring, whereas no changes were observed in the OFC (**Fig. 6j**). mHFD had no effect on serotonergic synapse density in either the NAc or OFC of adult female offspring (**Fig. 6k**) consistent with the sex and circuit specific changes observed in this model. To determine if the increased serotonergic synapse density in mHFD offspring resulted in enhanced serotonin release in the NAc, we measured serotonin release using the genetically encoded serotonin sensor (GRAB 5HT3.0)^56^. serotonergic fibers from *ex vivo* slices were electrically stimulated to evoke synchronized transmitter release, and stimulus-evoked fluorescence responses were imaged using two-photon microscopy (**Fig. 6l**). Electrical stimulation elicited robust GRAB 5HT3.0 fluorescence signals that scaled with stimulation frequency (**Fig. 6m**). Using this approach, we found that adult mHFD male offspring exhibited greater serotonin release in the NAc compared to mLFD male offspring (**Fig. 6n**), consistent with the hyperinnervation observed in this region (**Fig. 6b**). In contrast, mHFD did not alter serotonin release in the NAc of adult female offspring (**Fig. 6o**).

Serotonin functions as a critical developmental signaling molecule^90^, and elevated brain serotonin levels during sensitive periods (e.g., early life SSRI exposure) disrupts barrel-field organization^91–93^ and interneuron migration^94^. We therefore asked whether the serotonergic hyperinnervation observed in the NAc of P14 male offspring resulted in lasting alterations to excitatory or inhibitory synapse density in the adult NAc. To assess this, we quantified synapse density by measuring colocalization of synaptophysin with the glutamatergic postsynaptic maker PSD95 or the GABAergic postsynaptic marker gephyrin (**Extended data Fig. 3m**). We observed no effect of mHFD on either glutamatergic of GABAergic synapse density in the NAc of adult male offspring (**Extended data Fig. 3n,o**), suggesting that the effect of mHFD is restricted to the 5-HT system.

Given the gain in DRN→NAc circuit function and the increased 5-HT release into the NAc in adulthood, we determined whether mHFD affects adult behaviors mediated by the NAc or the OFC. To address this, we designed an operant task that assays both initial acquisition of lever pressing behavior, which is mediated in part by the NAc^95^, and reversal learning, a form of cognitive flexibility that depends on OFC function and is sensitive to serotonin manipulations^96–100^ (**Fig. 7a**). Because this task requires food restriction, we first assessed adult body weight to ensure that metabolic differences would not confound behavioral performance. Although mHFD offspring exhibited increased body weight during adolescence (P14-P21; **Extended data Fig.1a,b**), this difference was no longer evident in adulthood (2-4months) in either male or female offspring (**Extended data Fig.7a,b**) consistent with our previous results^11^. Importantly, no differences in body weight were observed between mHFD and mLFD offspring at either baseline or following food restriction (**Extended data Fig.7a,b**), indicating comparable metabolic status during behavioral testing.

**Figure 7.**
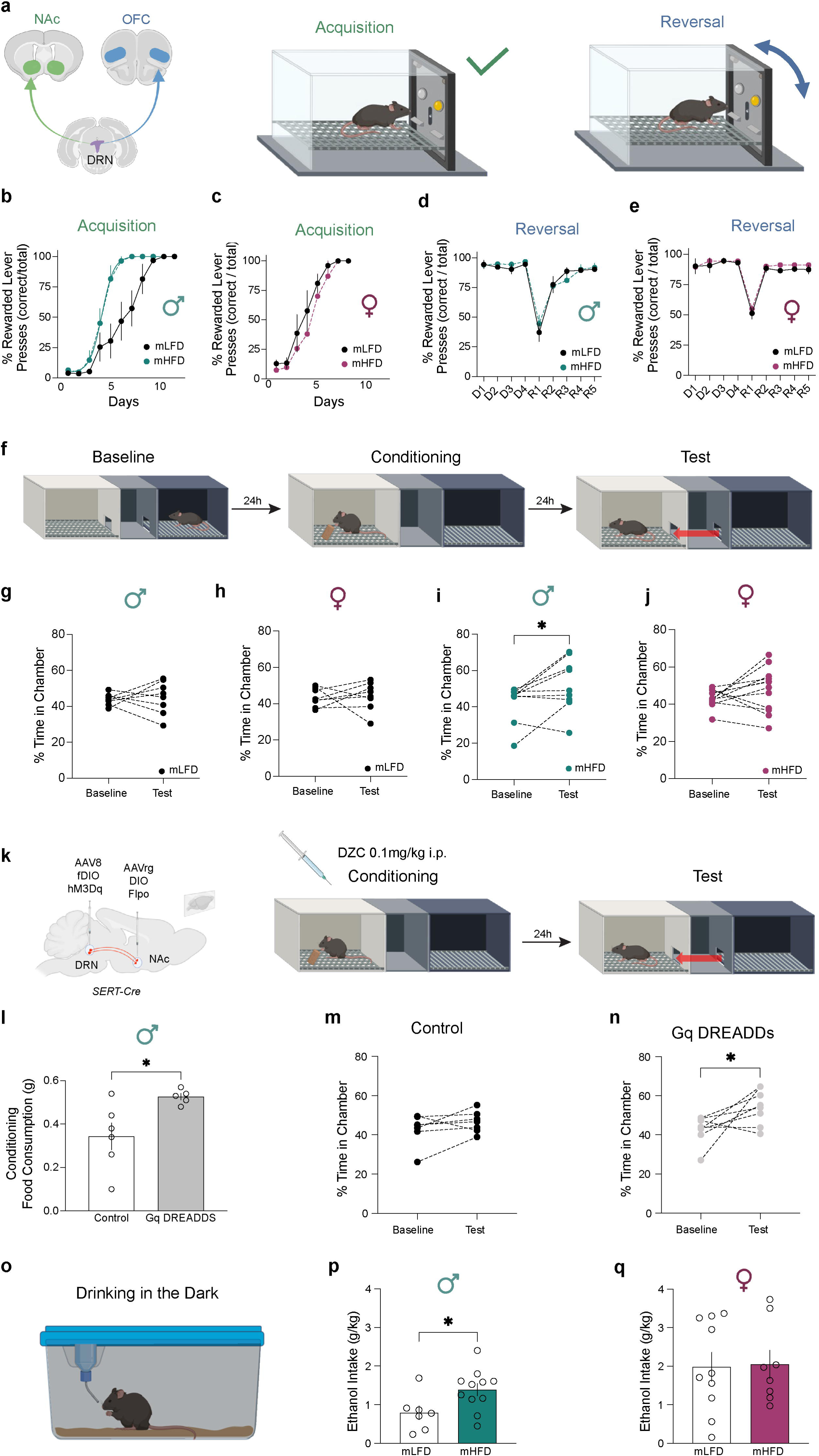
Maternal high-fat diet enhances reward-motivated learning in adult male but not female offspring. **a**, Schematic of operant task used to assess lever-press acquisition, which is mediated in part by the NAc, and reversal learning, which is mediated in part by the OFC. **b,c,** Learning curves for fixed-ratio 1 (FR1) lever-press acquisition in adult male (**b**) and female (**c**) offspring. **d,e**, Performance during single-discrimination training (D1-4) and reversal learning (R1-5) in adult male (**d**) and female (**e**) offspring. **f**, Schematic of the food-conditioned place preference (CPP) paradigm. **g**, mLFD male offspring do not exhibit a food-conditioned place preference following a single day of food conditioning (n=8; p=0.80; paired t-test). **h**, mLFD female offspring do not exhibit a food-conditioned place preference following a single day of food conditioning (n=8; p=0.80; paired t-test).**i**, In contrast, mHFD male offspring do exhibit a food-conditioned place preference following a single day of food conditioning (n=9; p<0.05; paired t-test), **j**, an effect no observed in mHFD female offspring (n=12; p=0.20; paired t-test). **k**, Schematic illustrating viral expression of Gq DREADDs in DRN→NAc serotonergic neurons and chemogentic activation of this circuit during food conditioning using Deschloroclozapine (DCZ). **l,** Chemogenetic activation of DRN→NAc serotonergic neurons increases food consumption during food conditioning session (n=6 control, 5 Gq DREADDs mLFD male offspring; p<0.05; Student’s t-test). **m**,mCherry-expressing control mLFD male offspring do not exhibit a food-conditioned place preference following a single food conditioning session (n=7; p=0.22; paired t-test). **n**, Chemogenetic activation of DRN→NAc serotonergic neurons during food conditioning is sufficient to induce a food-conditioned place preference in mLFD male offspring (n=8; p <0.05; paired t-test). **o**, Schematic of the Drinking in the Dark (DiD) paradigm. **p**, mHFD adult male offspring consume more ethanol than mLFD male offspring during the first day of DiD (n=7 mLFD, 11 mHFD male offspring; p <0.05; student’s t-test). **q**, mHFD adult female offspring do not consume more ethanol than mLFD offspring during the first day of DiD (n=10 mLFD, 19 mHFD male offspring; p =0.91; student’s t-test). Data represented as mean ± SEM. Gray points denote individual animals. *p<0.05.

During acquisition of the fixed-ratio 1 (FR1) lever pressing task, mHFD male offspring learned significantly faster than mLFD males, as evidenced by a leftward shift in the acquisition curve (**Fig. 7b**). This effect was not observed in female offspring (**Fig. 7c**). Consistent with this sex-specific enhancement, analysis of lever-pressing performance on day 6 of acquisition revealed a significant diet x sex interaction (**Extended data Fig.7c**), with mHFD male offspring achieving near mastery of FR1 training (mean = 96.3% correct lever presses) compared to mLFD male offspring (mean = 46.8% correct lever presses). No effect of mHFD on FR1 acquisition was detected in female offspring.

Following mastery of FR1 acquisition and subsequent single-discrimination training (defined as >80% correct performance for more than three consecutive days), mice underwent reversal learning in which the rewarded and non-rewarded levers were switched to assess updating of reward contingencies, a cognitive task that engages the OFC^96, 101–104^. Consistent with the absence of serotonergic remodeling in the OFC, mHFD did not affect reversal learning performance in either adult male or female offspring (**Fig. 7d-e**, **Extended data Fig.7d**).

Given the leftward shift in the FR1 acquisition curve observed in mHFD male offspring, we next asked whether these mice exhibit enhanced reward-motivated learning mediated by the NAc^95, 105^. To test this, we assessed reward learning using a conditioned-place preference (CPP) paradigm, a well-established assay of NAc-dependent learning^106–108^. We adapted a previously validated food CPP protocol in C57BL/6J mice where three days of conditioning are required to elicit a robust food-associated place preference. Since mHFD male offspring exhibited accelerated learning in the operant task (**Fig. 7b**), we employed an abbreviated CPP paradigm consisting of a single day of food conditioning (**Fig. 7f**). As expected, a single conditioning session was insufficient to induce a place preference in mLFD male or female offspring (**Fig. 7g,h**), indicating a subthreshold learning experience. However, mHFD male offspring exhibited a significant food-conditioned place preference (**Fig. 7i**), consistent with enhanced reward learning. This effect was not observed in mHFD female offspring (**Fig. 7j**), paralleling the sex-specific effects seen in operant learning and serotonin fiber innervation.

To determine if increased activity of the DRN→NAc serotonin circuit is sufficient to drive the enhanced food-motivated learning observed in mHFD male offspring, we used an intersectional viral strategy to express the excitatory DREADD receptor (hM3Dq) or mCherry control, selectively in DRN→NAc serotonin neurons (**Fig. 7k**). DREADDS expression was restricted to TPH2 positive neurons in the DRN (**Extended data Fig. 7e**), and administration of the selective DREADD agonist deschloroclozapine (DCZ) increased the firing rate in DRN→NAc serotonin neurons (**Extended data Fig. 7f**). Because DRN neurons increase their firing rate in response to rewarding and appetitive stimuli^109–112^, we activated the DRN→NAc circuit during the single food conditioning session (**Fig. 7k**). Mice received a vehicle injection (0.5% DMSO in saline, i.p.) during conditioning to the non-food paired chamber. Chemogenetic activation of DRN→NAc serotonin neurons increased food consumption during the conditioning session compared to mCherry control (**Fig. 7l**). During testing, mCherry-expressing control mLFD male offspring did not exhibit a food-conditioned place preference (**Fig. 7m)** consistent with prior results (**Fig. 7g**). In contrast, chemogenetic activation of DRN→NAc serotonin neurons during food conditioning was sufficient to induce a food-conditioned place preference in mLFD male offspring (**Fig. 7n**), phenocopying what was observed in mHFD male offspring. However, in mLFD female offspring, we did not observe an increase in food consumption during the single food conditioning session (**Extended data Fig. 7g,h**), but we did observe a significant food-conditioned place preference in both control (**Extended data Fig. 7i**) and Gq DREADDs treated mice (**Extended data Fig. 7j**).

To determine whether the effects of mHFD on reward-motivated behavior extend beyond food motivated behavior to broader reward processing, including behaviors directed toward drugs of abuse, we assessed voluntary ethanol consumption in adult offspring, as C57BL/6J mice readily consume 20% ethanol^71^. We used the drinking in the dark (DiD) paradigm, a well-established assay of voluntary binge-like ethanol consumption^70^ (**Fig. 7o**) and abbreviated the protocol to one week to specifically capture engagement of the NAc circuitry during early stage of ethanol consumption^113–115^. Because the DiD paradigm requires switching mice to a reverse light cycle, we first evaluated whether mHFD affected circadian entrainment, given the known role of serotonergic signaling in light-dependent circadian regulation^116, 117^. In a separate cohort of adult male mice, we recorded 24h wheel-running activity across a standard 12:12h light-dark cycle, constant darkness, and during acute light re-entrainment (**Extended data Fig. 7k,l**). mHFD did not alter overall locomotor activity during either the light or dark phase (**Extended data Fig. 7m**), nor did it affect circadian phase shift during free-running conditions (**Extended data Fig. 7n**) or in response to a 4h light pulse (**Extended data Fig. 7o**), indicating intact circadian entrainment.

Using the DiD paradigm, we found that adult mHFD male offspring consumed more ethanol than mLFD controls on the first day of access (**Fig. 7p**), an effect that normalized over subsequent drinking sessions (**Extended data Fig. 7p,q**). This effect was not observed in adult female offspring (**Fig. 7q; Extended data Fig. 7r**).

## Discussion

Maternal metabolic state is increasingly recognized as a determinate of offspring neurodevelopmental risk^6, 118–121^. In the U.S., more than 50% of women enter pregnancy overweight or obese^122, 123^, highlighting maternal metabolic state as a prevalent environmental factor likely contributing to the rising incidence of neurodevelopmental disorders^124–128 129–132^. Here we show that maternal high-fat diet (mHFD) exposure selectively disrupts postnatal development of a projection-defined serotonergic circuit in a sex-specific manner. In male offspring, dorsal raphe to nucleus accumbens (DRN→NAc) serotonergic projections exhibit postnatal hyperinnervation that persists into adulthood, whereas DRN to orbitofrontal cortex (DRN→OFC) projections remain largely unaffected. This projection-selective vulnerability reveals that serotonergic networks differ in developmental sensitivity to maternal diet and identifies reward circuitry as a preferential target of metabolic perturbation.

Our findings identify impaired microglial pruning as a candidate mechanism underlying this circuit-specific remodeling. Hyperinnervation of serotoninergic fibers in the NAc coincided with reduced microglial engulfment of serotonergic material during a critical postnatal window. Microglia are central regulators of axonal refinement and synapse elimination^35, 38, 133, 134^, and serotonergic signaling is known to modulate microglia phagocytic machinery^80, 83^. Our data extend this framework by implicating microglia-dependent pruning of neuromodulator projections as a diet-sensitive developmental process. We further suggest that the resulting circuit selective remodeling arises, at least in part, from regional heterogeneity in microglial 5-HT receptor expression. Using a novel viral strategy, we found that overexpression of the 5-HT2C receptor in NAc microglia, phenocopying what is observed in mHFD offspring, was sufficient to induce hyper serotonergic innervation. These results support a role for microglial serotonergic signaling in shaping the development of the 5-HT system.

In parallel, we examined whether the selective sensitivity of this circuit to mHFD was attributable to differences within the DRN. DRN→NAc and DRN→OFC neurons were largely non-overlapping populations with distinct transcriptional and functional signatures. However, we observed neither dorsoventral spatial segregation nor *Vglut3*-based segregation between these projection-defined populations, despite previous reports of spatial and molecular differences between cortical and subcortical projecting DRN neurons^73, 74, 88, 89^. We observed selective physiological remodeling of DRN→NAc serotonergic neurons in male mHFD offspring. These neurons exhibited reduced tonic firing, which may represent a homeostatic adaptation to increased terminal field density and transmitter release capacity. By contrast, their firing response during depolarizing drive was preserved, indicating maintained phasic excitability. Additionally, we observed reduced inhibitory gating of these neurons. Taken together, these data demonstrate that mHFD alters the DRN→NAc circuit at both the level of the DRN and its terminal field. These adaptations are predicted to collectively enhance serotonin output during behaviorally relevant circuit recruitment. Consistent with this interpretation, adult mHFD male offspring showed increased stimulus-evoked serotonin release in the NAc.

At the behavioral level, this circuit remodeling was associated with augmented reward-motivated learning. The effect was restricted to the acquisition phase rather than sustained reward consumption as adult body weights did not differ across diet groups. Chemogenetic activation of DRN→NAc serotonergic neurons phenocopied the enhanced reward-learning phenotype observed in mHFD male offspring, establishing circuit-level sufficiency. Serotonin release in the NAc regulates cortico-striatal plasticity^135^ and local dopamine release^136–138^, providing plausible mechanisms through which elevated serotonergic signaling could amplify reward learning and motivational vigor. Extension of this phenotype from food reward to alcohol suggests that maternal diet may broadly tune mesolimbic circuit gain rather than affect a single behavioral domain.

The male specificity of these effects parallels the sex bias observed in several neurodevelopmental disorders. Sex differences in microglial maturation^139, 140^, immune signaling^141, 142^, and serotonergic development may jointly contribute to this vulnerability, although the precise mechanisms remain to be determined. Notably, viral upregulation of the 5-HT2C receptor increased 5-HT fiber density in both male and female offspring. Likewise, mHFD increased 5-HT2C receptor expression in both sexes. However, only male mHFD offspring exhibited serotonergic hyperinnervation, indicating that receptor upregulation alone is insufficient to account for the sex-specific phenotype. These findings suggest that an additional sex-dependent mechanism operates in the context of mHFD. One possibility is sex differences in immune signaling interact with elevated microglial 5-HT2C receptor expression to alter serotonergic pruning. Such signaling may be engaged in both sexes following stereotaxic surgery to account for hyperinnervation observed in both sexes following viral receptor upregulation. Future studies dissecting the contribution of inflammatory signaling will be important for solving sex-specific vulnerability to mHFD. In summary, our study identifies a projection-specific mechanism by which maternal diet reshapes serotonergic reward circuity. More broadly, this work demonstrates that neuromodulator systems exhibit circuit-level heterogeneity in developmental resilience, which underscores the importance of projection-resolved analysis for understanding and ultimately targeting serotonergic dysfunction in neurodevelopmental disorders.

---

**Extended Data Figure 1. Maternal high-fat diet does not alter serotonergic innervation in the OFC at P14 or the NAc at P21**. **a,** mHFD increases body weight in male and female offspring at P14 (n=13 males, 19 females, p<0.01; two-way ANOVA). **b**, This increase in body weight persists at P21 (n=26 males, 17 females; p<0.01; two-way ANOVA ). **c-f**, mHFD does not alter serotonergic innervation of the OFC at P14 in male (n=6 mLFD, 6 mHFD; p=0.63; Student’s t- test) or female offspring (n=6 mLFD, 6 mHFD; p=0.17; Student’s t-test). **g-j**, mHFD-induced hyperinnervation of the NAc is no longer evident at P21 in male offspring (n=6 mLFD, 6 mHFD; p=0.29; Student’s t-test) and is still not observed in female offspring at this age (n=6 mLFD, 6 mHFD; p=0.18; Student’s t-test). Data represented as mean ± SEM. Gray points denote individual images. *p<0.05, **p<0.01.

**Extended Data Figure 2. Maternal high-fat diet does not impact microglial phagocytosis of serotonergic fibers in female offspring**. **a,** Representative 3D reconstructions of NAc microglia from mLFD and mHFD female offspring at P14, showing lysosomal compartment (CD68, red) containing serotonergic material (SERT, green). Scale bar 7µm. **b**, NAc microglia from mHFD female offspring show a trend toward reduced lysosomal volume compared with mLFD offspring at P14 (p=0.05; Student’s t-test). **c**, No difference in SERT phagocytosis is observed in NAc microglia at P14 (n=6 mLFD, 6mHFD offspring; p=0.86; Student’s t-test). **d**, Representative 3D reconstructions of NAc microglia from mLFD and mHFD female offspring at P21. **e,f,** mHFD does not alter lysosomal volume (**e**, p=0.80) or SERT phagocytosis (**f**, p=0.15) in NAc microglia at P21 (n=6 mLFD, 6mHFD offspring; Student’s t-test). **g**, Representative 3D reconstructions of OFC microglia from mLFD and mHFD female offspring at P14, showing CD68-positive lysosomes (red) containing SERT (green). Scale bar 7µm. **h**,**i,** mHFD does not alter lysosomal volume (**h**, p=0.43) or SERT phagocytosis (**i**, p=0.59) in OFC microglia at P14 (n=6 mLFD, 6mHFD offspring; Student’s t-test). **j**, Representative 3D reconstructions of OFC microglia from mLFD and mHFD female offspring at P21. **k**, OFC microglia from mHFD female offspring exhibit increased lysosomal volume compared with mLFD offspring at P21 (n=6 mLFD, 6mHFD offspring; p<0.01; Student’s t-test). **l**, mHFD does not affect SERT phagocytosis in OFC microglia at P21 (n=6 mLFD, 6mHFD offspring; p=0.29; Student’s t-test). Data represented as mean ± SEM. Gray points denote individual cells. # p=0.05, **p<0.01.

**Extended Data Figure 3. Serotonin receptor heterogeneity across the nucleus accumbens and orbitofrontal cortex**. **a**, Non-microglial (CD11B negative) cells isolated from the NAc of P14 offspring express higher *Htr1b* (p<0.01), *Htr2b* (p<0.01), and *Htr2c* (p<0.01) compared with cells from the OFC (n=8 males, 8 females per brain region; Student’s t-test). In contrast, NAc non- microglial cells exhibit lower expression of *Htr1a* (p<0.01) and *Htr2a* (p<0.01) relative to OFC cells. **b**, In adult (2-4 months) microglia, expression of *Htr1a* (p<0.01) and *Htr2a* (p<0.01) is higher in the OFC, whereas *Htr2c* expression is higher in the NAc (p<0.01)(n=4 males, 4 females per brain region; Student’s t-test). No regional differences were observed in expression of *Htr1b* (p=0.56) or *Htr2b* (p=0.73). **c**, In adult non-microglia (Cd11b-negative) cells, expression of *Htr1a* (p<0.05) and *Htr2a* (p<0.05) is higher in the OFC, whereas expression of *Htr1b* (p<0.01) and *Htr2c* (p<0.01) is higher in the NAc, with a trend toward increased 5HTR-2B expression (p=0.06) (n=4 males, 4 females per brain region; Student’s t-test). **d**, In non-microglial (CD11B negative) cells isolated from the NAc of P14 offspring, mHFD does not affect expression of *Htr1a* (p=0.86), *Htr1b* (p=0.35), *Htr2a* (p=0.49), or *Htr2b* (p=0.93). A non-significant trend toward increased expression of *Htr2c* is observed (p=0.11) (n= 7 males,7 females mLFD; 7 males 6 females mHFD; Student’s t-test). Data represented as mean ± SEM. *p<0.05, **p<0.01.

**Extended Data Figure 4. Microglia-targeting AAV to genetically manipulate nucleus accumbens development. a**, Representative 60x confocal images of AAV-GFP in IBA1 positive microglia. **b**, AAV-GFP expression is largely restricted to IBA1 positive microglia with little to no expression evident in IBA1 negative cells (n= 5 males, 4 females). **c**, Representative schematic of AAV plasmid expressing the 5-HT2C receptor under the mouse IBA1 promoter. **d**, Representative 3D reconstructions of microglia (IBA1) and Htr2c puncta from RNA-FISH. e, NAc microglia from mLFD P14 offspring treated with the AAV plasmid expressing the 5-HT2C receptor exhibit greater volume of Htr2c puncta compared to GFP-control treated offspring (p <0.05; n= 3 males, 3 females per group; Student’s t-test). Data represented as mean ± SEM. *p<0.05

**Extended Data Figure 5. DRN→OFC serotonergic neurons do not preferentially express cortical marker Vglut3. a,** Schematic depicting intersectional viral strategy to label projection- defined serotonergic DRN neurons. **b**, Representative confocal image depicting serotonin neurons (TPH2 = green) and AAV mCherry (red) in the DRN. AAV mCherry signal overlaps with TPH2 positive serotonin neurons. **c**, Schematic depicting microinjection of a retrograde AAV expressing cre recombinase by the TPH2 promoter in Ribotag mice. **d**, Representative confocal images of the DRN depicting TPH2 (green) and viral cre recombinase (magenta). **e-g,** dot plots depicting top differentially expressed genes across the five serotonin neuron subtypes classified from the single nuclear sequencing study clustered by (**e**) channels, (**f**) neurotransmission, and (**g**) transcription factors. OFC projecting neurons express greater Trpc3, En2, Pou3f3, Nr3c1 but less Pdyn, Thr, and Nr2f2 compared to NAc projecting neurons. There was no difference in resting membrane potential (**h**; p=0.72) or action potential threshold (**i**; p=0.17) between DRN→NAc and DRN→OFC serotonergic neurons (student’s t-test). **j**, Representative RNA-FISH images showing *Vglut3* (magenta), *Gad2* (green), and viral mCherry (red) in the DRN. Scale bars = 100µm, 20µm. **k**, Quantification of *Vglut3* and *Gad2* expression in DRN→NAc and DRN→OFC serotonergic neuron shows no difference between projection-defined subtype. Data represented as mean ± SEM.

**Extended Data Figure 6. Maternal high-fat diet does not alter glutamatergic or GABAergic synapse density in the nucleus accumbens**. **a**, Representative trace showing action potential firing in response to depolarizing current injection in a DRN→NAc serotonergic neuron. mHFD does not alter firing response to depolarizing current injection in DRN→NAc (**b**, p=0.33) or DRN→OFC (**c**, p=0.37) serotonergic neurons from female offspring (n=6 mLFD, 6mHFD offspring per group; two-way repeated measures ANOVA). **d**, Representative trace showing membrane potential response to ramping current injection (200pA s^−1^). mHFD does not alter action potential threshold in DRN→NAc (**e**, p=0.78) or DRN→OFC (**f**, p=0.33) serotonergic neurons (NAc: n= 7 mLFD males, 6 mLFD females, 6 mHFD males, 6 mHFD females; OFC: n=6 mLFD males, 6 mLFD females, 6 mHFD males, 6 mHFD females; two-way ANOVA). mHFD does not alter input resistance in DRN→NAc (**g**, p=0.82) or DRN→OFC (**h**, p=0.79) serotonergic neurons (same n as in **e,f**; two-way ANOVA). **i**, Sholl analysis reveals no impact of mHFD on DRN→NAc serotonergic neuron neurite morphology in adult male offspring (n=16mLFD, 15mHFD cells; p=0.76; Mixed- effects model). **j**, Representative miniature excitatory post-synaptic current (mEPSC) traces from mLFD (black) and mHFD (teal) DRN→NAc serotonergic neurons. scale bar = 60pA, 15s. mHFD induces a non-significant, but trending increase in mEPSC frequency (**k**; p=0.12), but no change in amplitude (**l**; p=0.41) on DRN→NAc serotonergic neurons (n=5 mLFD, 4 mHFD male offspring; Student’s t-test). **m**, Representative 60x confocal images of synaptophysin (green), the glutamatergic postsynaptic marker PSD95 (red) and the GABAergic postsynaptic marker gephyrin (purple) in the NAc of adult male offspring. Scale bar 20µm. mHFD does not alter glutamatergic (**n**, p=0.48) or GABAergic (**o**, p=0.79) synapse density in the NAc of adult male offspring (n=6 mLFD, 6 mHFD male offspring; Student’s t-test). Data represented as mean ± SEM. Gray points denote individual cells or images.

**Extended Data Figure 7. Maternal high-fat diet does not alter body weight or circadian behavior in adult offspring**. **a**, mHFD does not affect body weight at baseline or following food restriction in adult (2-4 months) male offspring (n=8 mLFD, 8 mHFD; p=0.49; two-way repeated measures ANOVA). **b**, mHFD does not affect body weight at baseline or following food restriction in adult (2-4 months) female offspring (n=6 mLFD, 7 mHFD; p=0.83; two-way repeated measures ANOVA).**c**, mHFD male offspring acquire the fixed-ratio 1 (FR1) lever-pressing task faster than mLFD males, whereas no effect of diet is observed in females (n=7 male + 6 female mLFD, 7 male + 7 female mHFD; Interaction: p <0.05; two-way ANOVA; Sidak’s multiple comparisons test p>0.01). **d**, mHFD does not affect day 1 reversal learning performance (main effect of diet: p=0.36; two-way ANOVA). **e**, Representative confocal image showing Gq DREADDS expression (red) in TPH2-positive serotonergic neurons (green) in the DRN. Scale bars, 100µm (left) and 10µm (right). **f**, Representative current-clamp recording showing membrane depolarization and action potential firing in response to bath application of deschloroclozapine (DCZ;10µM). **g**, Schematic illustrating viral expression of Gq DREADDs in DRN→NAc serotonergic neurons. **h,** Chemogenetic activation of DRN→NAc serotonergic neurons does not increase food consumption during food conditioning session in female mLFD offspring (n= 8 control, 7 Gq DREADDs mLFD offspring; p=0.93; Student’s t-test). **i**,mCherry-expressing control mLFD female offspring exhibit a food-conditioned place preference following a single food conditioning session (n=8; p<0.05; paired t-test). **j**, a food-conditioned place preference was also observed in mLFD female offspring expressing Gq DREADDs in DRN→NAc serotonergic neurons (n=7; p <0.01; paired t-test). **k**, Schematic of the running-wheel assay during 12:12-h light-dark cycle and during constant darkness (free run). **l**, Representative actograms from adult mLFD and mHFD male offspring during 12:12-h light entrainment, 24-h constant darkness (free run), a 4-h light pulse (asterisk), and re-entrainment to 12:12-h light-dark cycle. **m**, mHFD does not impact running wheel activity during light (n=4 mLFD, 4 mHFD males; p=0.41; two-way repeated measures ANOVA or dark phase of the 12:12h entrainment period (p=0.89; two-way repeated measures ANOVA). **n**, mHFD does not impact phase shift during 24h free run (p=0.84; two-way repeated measures ANOVA). **o**, mHFD does not impact phase shift in response to 4h light pulse (p=0.56; Student’s t-test). **p**, Schematic of the Drinking in the Dark (DiD) paradigm. **q**, mHFD increases initial ethanol consumption in adult male offspring that normalizes across 1-week of voluntary drinking (n=7 mLFD, 11 mHFD male offspring; time x diet interaction: p <0.05; Two-way repeated measures ANOVA). **r**, mHFD does not impact adult female ethanol consumption (n=10 mLFD, 10 mHFD female offspring; main effect of diet: p =0.93; Two-way repeated measures ANOVA). Data represented as mean ± SEM. Gray points denote individual animals. **p<0.01.

## References

1. Khaire A, Wadhwani N, Madiwale S, Joshi S. Maternal fats and pregnancy complications: Implications for long-term health. Prostaglandins Leukot Essent Fatty Acids. 2020;157:102098. Epub 20200421. doi: 10.1016/j.plefa.2020.102098. PubMed PMID: 32380367.

2. Lau C, Rogers JM, Desai M, Ross MG. Fetal programming of adult disease: implications for prenatal care. Obstet Gynecol. 2011;117(4):978–85. doi: 10.1097/AOG.0b013e318212140e. PubMed PMID: 21422872.

3. Bordeleau M, Fernández de Cossío L, Chakravarty MM, Tremblay M. From Maternal Diet to Neurodevelopmental Disorders: A Story of Neuroinflammation. Front Cell Neurosci. 2020;14:612705. Epub 20210115. doi: 10.3389/fncel.2020.612705. PubMed PMID: 33536875; PMCID: PMC7849357.

4. Ferreira LB, Lobo CV, Miranda A, Carvalho BDC, Santos LCD. Dietary Patterns during Pregnancy and Gestational Weight Gain: A Systematic Review. Rev Bras Ginecol Obstet. 2022;44(5):540–7. Epub 20220428. doi: 10.1055/s-0042-1744290. PubMed PMID: 35483873; PMCID: PMC9948295.

5. Cummings JR, Lipsky LM, Schwedhelm C, Liu A, Nansel TR. Associations of ultra-processed food intake with maternal weight change and cardiometabolic health and infant growth. Int J Behav Nutr Phys Act. 2022;19(1):61. Epub 20220526. doi: 10.1186/s12966-022-01298-w. PubMed PMID: 35619114; PMCID: PMC9137185.

6. Getz KD, Anderka MT, Werler MM, Jick SS. Maternal Pre-pregnancy Body Mass Index and Autism Spectrum Disorder among Offspring: A Population-Based Case-Control Study. Paediatr Perinat Epidemiol. 2016;30(5):479–87. Epub 20160530. doi: 10.1111/ppe.12306. PubMed PMID: 27239935; PMCID: PMC5849232.

7. Wang Y, Tang S, Xu S, Weng S, Liu Z. Maternal Body Mass Index and Risk of Autism Spectrum Disorders in Offspring: A Meta-analysis. Sci Rep. 2016;6:34248. Epub 20160930. doi: 10.1038/srep34248. PubMed PMID: 27687989; PMCID: PMC5043237.

8. Li L, Lagerberg T, Chang Z, Cortese S, Rosenqvist MA, Almqvist C, D’Onofrio BM, Hegvik TA, Hartman C, Chen Q, Larsson H. Maternal pre-pregnancy overweight/obesity and the risk of attention-deficit/hyperactivity disorder in offspring: a systematic review, meta-analysis and quasi-experimental family-based study. Int J Epidemiol. 2020;49(3):857–75. doi: 10.1093/ije/dyaa040. PubMed PMID: 32337582; PMCID: PMC7394963.

9. Kacperska M, Mizera J, Pilecki M, Pomierny-Chamioło L. The impact of excessive maternal weight on the risk of neuropsychiatric disorders in offspring-a narrative review of clinical studies. Pharmacol Rep. 2024;76(3):452–62. Epub 20240422. doi: 10.1007/s43440-024-00598-1. PubMed PMID: 38649593; PMCID: PMC11126479.

10. Frazier JA, Li X, Kong X, Hooper SR, Joseph RM, Cochran DM, Kim S, Fry RC, Brennan PA, Msall ME, Fichorova RN, Hertz-Picciotto I, Daniels JL, Lai JS, Boles RE, Zvara BJ, Jalnapurkar I, Schweitzer JB, Singh R, Posner J, Bennett DH, Kuban KCK, O’Shea TM. Perinatal Factors and Emotional, Cognitive, and Behavioral Dysregulation in Childhood and Adolescence. J Am Acad Child Adolesc Psychiatry. 2023;62(12):1351–62. Epub 20230517. doi: 10.1016/j.jaac.2023.05.010. PubMed PMID: 37207889; PMCID: PMC10654259.

11. Ceasrine AM, Devlin BA, Bolton JL, Green LA, Jo YC, Huynh C, Patrick B, Washington K, Sanchez CL, Joo F, Campos-Salazar AB, Lockshin ER, Kuhn C, Murphy SK, Simmons LA, Bilbo SD. Maternal diet disrupts the placenta-brain axis in a sex-specific manner. Nat Metab. 2022;4(12):1732–45. Epub 20221128. doi: 10.1038/s42255-022-00693-8. PubMed PMID: 36443520; PMCID: PMC10507630.

12. Fernandes DJ, Spring S, Roy AR, Qiu LR, Yee Y, Nieman BJ, Lerch JP, Palmert MR. Exposure to maternal high-fat diet induces extensive changes in the brain of adult offspring. Transl Psychiatry. 2021;11(1):149. Epub 20210302. doi: 10.1038/s41398-021-01274-1. PubMed PMID: 33654064; PMCID: PMC7925669.

13. Kang SS, Kurti A, Fair DA, Fryer JD. Dietary intervention rescues maternal obesity induced behavior deficits and neuroinflammation in offspring. J Neuroinflammation. 2014;11:156. Epub 20140912. doi: 10.1186/s12974-014-0156-9. PubMed PMID: 25212412; PMCID: PMC4172780.

14. Bordeleau M, Comin CH, Fernández de Cossío L, Lacabanne C, Freitas-Andrade M, González Ibáñez F, Raman-Nair J, Wakem M, Chakravarty M, Costa LDF, Lacoste B, Tremblay M. Maternal high-fat diet in mice induces cerebrovascular, microglial and long-term behavioural alterations in offspring. Commun Biol. 2022;5(1):26. Epub 20220111. doi: 10.1038/s42003-021-02947-9. PubMed PMID: 35017640; PMCID: PMC8752761.

15. Ma J, Prince AL, Bader D, Hu M, Ganu R, Baquero K, Blundell P, Alan Harris R, Frias AE, Grove KL, Aagaard KM. High-fat maternal diet during pregnancy persistently alters the offspring microbiome in a primate model. Nat Commun. 2014;5:3889. Epub 20140520. doi: 10.1038/ncomms4889. PubMed PMID: 24846660; PMCID: PMC4078997.

16. Rivera HM, Kievit P, Kirigiti MA, Bauman LA, Baquero K, Blundell P, Dean TA, Valleau JC, Takahashi DL, Frazee T, Douville L, Majer J, Smith MS, Grove KL, Sullivan EL. Maternal high-fat diet and obesity impact palatable food intake and dopamine signaling in nonhuman primate offspring. Obesity (Silver Spring). 2015;23(11):2157–64. doi: 10.1002/oby.21306. PubMed PMID: 26530932; PMCID: PMC4636015.

17. True C, Dean T, Takahashi D, Sullivan E, Kievit P. Maternal High-Fat Diet Effects on Adaptations to Metabolic Challenges in Male and Female Juvenile Nonhuman Primates. Obesity (Silver Spring). 2018;26(9):1430–8. doi: 10.1002/oby.22249. PubMed PMID: 30226008; PMCID: PMC6146409.

18. Sun P, Wang M, Chai X, Liu YX, Li L, Zheng W, Chen S, Zhu X, Zhao S. Disruption of tryptophan metabolism by high-fat diet-triggered maternal immune activation promotes social behavioral deficits in male mice. Nat Commun. 2025;16(1):2105. Epub 20250302. doi: 10.1038/s41467-025-57414-4. PubMed PMID: 40025041; PMCID: PMC11873046.

19. Gohir W, Kennedy KM, Wallace JG, Saoi M, Bellissimo CJ, Britz-McKibbin P, Petrik JJ, Surette MG, Sloboda DM. High-fat diet intake modulates maternal intestinal adaptations to pregnancy and results in placental hypoxia, as well as altered fetal gut barrier proteins and immune markers. J Physiol. 2019;597(12):3029–51. Epub 20190513. doi: 10.1113/jp277353. PubMed PMID: 31081119.

20. Wallace JG, Bellissimo CJ, Yeo E, Fei Xia Y, Petrik JJ, Surette MG, Bowdish DME, Sloboda DM. Obesity during pregnancy results in maternal intestinal inflammation, placental hypoxia, and alters fetal glucose metabolism at mid-gestation. Sci Rep. 2019;9(1):17621. Epub 20191126. doi: 10.1038/s41598-019-54098-x. PubMed PMID: 31772245; PMCID: PMC6879619.

21. Arslan S, Yıldıran H, Seymen CM. The Effect of Maternal High-Fat Diet on Adipose Tissue Histology and Lipid Metabolism-Related Genes Expression in Offspring Rats. Nutrients. 2024;16(1). Epub 20240102. doi: 10.3390/nu16010150. PubMed PMID: 38201978; PMCID: PMC10780511.

22. Smoothy J, Larcombe AN, Chivers EK, Matthews VB, Gorman S. Correction to: Maternal high fat diet compromises survival and modulates lung development of offspring, and impairs lung function of dams (female mice). Respir Res. 2021;22(1):81. Epub 20210314. doi: 10.1186/s12931-021-01681-4. PubMed PMID: 33715622; PMCID: PMC7958411.

23. Woods RM, Lorusso JM, Fletcher J, ElTaher H, McEwan F, Harris I, Kowash HM, D’Souza SW, Harte M, Hager R, Glazier JD. Maternal immune activation and role of placenta in the prenatal programming of neurodevelopmental disorders. Neuronal Signal. 2023;7(2):Ns20220064. Epub 20230531. doi: 10.1042/ns20220064. PubMed PMID: 37332846; PMCID: PMC10273029.

24. Zaretsky MV, Alexander JM, Byrd W, Bawdon RE. Transfer of inflammatory cytokines across the placenta. Obstet Gynecol. 2004;103(3):546–50. doi: 10.1097/01.Aog.0000114980.40445.83. PubMed PMID: 14990420.

25. Sullivan EL, Riper KM, Lockard R, Valleau JC. Maternal high-fat diet programming of the neuroendocrine system and behavior. Horm Behav. 2015;76:153–61. Epub 20150424. doi: 10.1016/j.yhbeh.2015.04.008. PubMed PMID: 25913366; PMCID: PMC4619177.

26. Edlow AG. Maternal obesity and neurodevelopmental and psychiatric disorders in offspring. Prenat Diagn. 2017;37(1):95–110. Epub 20161107. doi: 10.1002/pd.4932. PubMed PMID: 27684946; PMCID: PMC5572633.

27. Shook LL, Kislal S, Edlow AG. Fetal brain and placental programming in maternal obesity: A review of human and animal model studies. Prenat Diagn. 2020;40(9):1126–37. Epub 20200517. doi: 10.1002/pd.5724. PubMed PMID: 32362000; PMCID: PMC7606714.

28. Harmancıoğlu B, Kabaran S. Maternal high fat diets: impacts on offspring obesity and epigenetic hypothalamic programming. Front Genet. 2023;14:1158089. Epub 20230511. doi: 10.3389/fgene.2023.1158089. PubMed PMID: 37252665; PMCID: PMC10211392.

29. Cunningham CL, Martínez-Cerdeño V, Noctor SC. Microglia regulate the number of neural precursor cells in the developing cerebral cortex. J Neurosci. 2013;33(10):4216–33. doi: 10.1523/jneurosci.3441-12.2013. PubMed PMID: 23467340; PMCID: PMC3711552.

30. Radecki DZ, Samanta J. Endogenous Neural Stem Cell Mediated Oligodendrogenesis in the Adult Mammalian Brain. Cells. 2022;11(13). Epub 20220702. doi: 10.3390/cells11132101. PubMed PMID: 35805185; PMCID: PMC9265817.

31. Reemst K, Noctor SC, Lucassen PJ, Hol EM. The Indispensable Roles of Microglia and Astrocytes during Brain Development. Front Hum Neurosci. 2016;10:566. Epub 20161108. doi: 10.3389/fnhum.2016.00566. PubMed PMID: 27877121; PMCID: PMC5099170.

32. Rigby MJ, Gomez TM, Puglielli L. Glial Cell-Axonal Growth Cone Interactions in Neurodevelopment and Regeneration. Front Neurosci. 2020;14:203. Epub 20200310. doi: 10.3389/fnins.2020.00203. PubMed PMID: 32210757; PMCID: PMC7076157.

33. Squarzoni P, Oller G, Hoeffel G, Pont-Lezica L, Rostaing P, Low D, Bessis A, Ginhoux F, Garel S. Microglia modulate wiring of the embryonic forebrain. Cell Rep. 2014;8(5):1271–9. Epub 20140821. doi: 10.1016/j.celrep.2014.07.042. PubMed PMID: 25159150.

34. Gesuita L, Cavaccini A, Argunsah A, Favuzzi E, Ibrahim LA, Stachniak TJ, De Gennaro M, Utz S, Greter M, Karayannis T. Microglia contribute to the postnatal development of cortical somatostatin-positive inhibitory cells and to whisker-evoked cortical activity. Cell Rep. 2022;40(7):111209. doi: 10.1016/j.celrep.2022.111209. PubMed PMID: 35977514; PMCID: PMC9396528.

35. Paolicelli RC, Bolasco G, Pagani F, Maggi L, Scianni M, Panzanelli P, Giustetto M, Ferreira TA, Guiducci E, Dumas L, Ragozzino D, Gross CT. Synaptic pruning by microglia is necessary for normal brain development. Science. 2011;333(6048):1456–8. Epub 20110721. doi: 10.1126/science.1202529. PubMed PMID: 21778362.

36. Hong S, Dissing-Olesen L, Stevens B. New insights on the role of microglia in synaptic pruning in health and disease. Curr Opin Neurobiol. 2016;36:128–34. Epub 20151230. doi: 10.1016/j.conb.2015.12.004. PubMed PMID: 26745839; PMCID: PMC5479435.

37. Scott-Hewitt N, Perrucci F, Morini R, Erreni M, Mahoney M, Witkowska A, Carey A, Faggiani E, Schuetz LT, Mason S, Tamborini M, Bizzotto M, Passoni L, Filipello F, Jahn R, Stevens B, Matteoli M. Local externalization of phosphatidylserine mediates developmental synaptic pruning by microglia. Embo j. 2020;39(16):e105380. Epub 20200713. doi: 10.15252/embj.2020105380. PubMed PMID: 32657463; PMCID: PMC7429741.

38. Schafer DP, Lehrman EK, Kautzman AG, Koyama R, Mardinly AR, Yamasaki R, Ransohoff RM, Greenberg ME, Barres BA, Stevens B. Microglia sculpt postnatal neural circuits in an activity and complement-dependent manner. Neuron. 2012;74(4):691–705. doi: 10.1016/j.neuron.2012.03.026. PubMed PMID: 22632727; PMCID: PMC3528177.

39. Roumier A, Pascual O, Béchade C, Wakselman S, Poncer JC, Réal E, Triller A, Bessis A. Prenatal activation of microglia induces delayed impairment of glutamatergic synaptic function. PLoS One. 2008;3(7):e2595. Epub 20080709. doi: 10.1371/journal.pone.0002595. PubMed PMID: 18612411; PMCID: PMC2440505.

40. Ikezu S, Yeh H, Delpech JC, Woodbury ME, Van Enoo AA, Ruan Z, Sivakumaran S, You Y, Holland C, Guillamon-Vivancos T, Yoshii-Kitahara A, Botros MB, Madore C, Chao PH, Desani A, Manimaran S, Kalavai SV, Johnson WE, Butovsky O, Medalla M, Luebke JI, Ikezu T. Inhibition of colony stimulating factor 1 receptor corrects maternal inflammation-induced microglial and synaptic dysfunction and behavioral abnormalities. Mol Psychiatry. 2021;26(6):1808–31. Epub 20200218. doi: 10.1038/s41380-020-0671-2. PubMed PMID: 32071385; PMCID: PMC7431382.

41. Block CL, Eroglu O, Mague SD, Smith CJ, Ceasrine AM, Sriworarat C, Blount C, Beben KA, Malacon KE, Ndubuizu N, Talbot A, Gallagher NM, Chan Jo Y, Nyangacha T, Carlson DE, Dzirasa K, Eroglu C, Bilbo SD. Prenatal environmental stressors impair postnatal microglia function and adult behavior in males. Cell Rep. 2022;40(5):111161. doi: 10.1016/j.celrep.2022.111161. PubMed PMID: 35926455; PMCID: PMC9438555.

42. Bilbo SD, Schwarz JM. Early-life programming of later-life brain and behavior: a critical role for the immune system. Front Behav Neurosci. 2009;3:14. Epub 20090824. doi: 10.3389/neuro.08.014.2009. PubMed PMID: 19738918; PMCID: PMC2737431.

43. Hanamsagar R, Bilbo SD. Environment matters: microglia function and dysfunction in a changing world. Curr Opin Neurobiol. 2017;47:146–55. Epub 20171106. doi: 10.1016/j.conb.2017.10.007. PubMed PMID: 29096243; PMCID: PMC5732848.

44. Morales L, Del Olmo N, Valladolid-Acebes I, Fole A, Cano V, Merino B, Stucchi P, Ruggieri D, López L, Alguacil LF, Ruiz-Gayo M. Shift of circadian feeding pattern by high-fat diets is coincident with reward deficits in obese mice. PLoS One. 2012;7(5):e36139. Epub 20120503. doi: 10.1371/journal.pone.0036139. PubMed PMID: 22570696; PMCID: PMC3343034.

45. Dunn GA, Thompson JR, Mitchell AJ, Papadakis S, Selby M, Fair D, Gustafsson HC, Sullivan EL. Perinatal Western-style diet alters serotonergic neurons in the macaque raphe nuclei. Front Neurosci. 2022;16:1067479. Epub 20230110. doi: 10.3389/fnins.2022.1067479. PubMed PMID: 36704012; PMCID: PMC9872117.

46. Esposito D, Cruciani G, Zaccaro L, Di Carlo E, Spitoni GF, Manti F, Carducci C, Fiori E, Leuzzi V, Pascucci T. A Systematic Review on Autism and Hyperserotonemia: State-of-the-Art, Limitations, and Future Directions. Brain Sci. 2024;14(5). Epub 20240510. doi: 10.3390/brainsci14050481. PubMed PMID: 38790459; PMCID: PMC11119126.

47. Chugani DC, Muzik O, Behen M, Rothermel R, Janisse JJ, Lee J, Chugani HT. Developmental changes in brain serotonin synthesis capacity in autistic and nonautistic children. Ann Neurol. 1999;45(3):287–95. doi: 10.1002/1531-8249(199903)45:3<287::aid-ana3>3.0.co;2-9. PubMed PMID: 10072042.

48. Beversdorf DQ, Nordgren RE, Bonab AA, Fischman AJ, Weise SB, Dougherty DD, Felopulos GJ, Zhou FC, Bauman ML. 5-HT2 receptor distribution shown by [18F] setoperone PET in high-functioning autistic adults. J Neuropsychiatry Clin Neurosci. 2012;24(2):191–7. doi: 10.1176/appi.neuropsych.11080202. PubMed PMID: 22772667.

49. Jackson EF, Riley TB, Overton PG. Serotonin dysfunction in ADHD. J Neurodev Disord. 2025;17(1):20. Epub 20250422. doi: 10.1186/s11689-025-09610-y. PubMed PMID: 40264019; PMCID: PMC12013068.

50. Seidl R, Kaehler ST, Prast H, Singewald N, Cairns N, Gratzer M, Lubec G. Serotonin (5-HT) in brains of adult patients with Down syndrome. J Neural Transm Suppl. 1999;57:221–32. doi: 10.1007/978-3-7091-6380-1_14. PubMed PMID: 10666678.

51. Gaspar P, Cases O, Maroteaux L. The developmental role of serotonin: news from mouse molecular genetics. Nat Rev Neurosci. 2003;4(12):1002–12. doi: 10.1038/nrn1256. PubMed PMID: 14618156.

52. Lidov HG, Molliver ME. An immunohistochemical study of serotonin neuron development in the rat: ascending pathways and terminal fields. Brain Res Bull. 1982;8(4):389–430. doi: 10.1016/0361-9230(82)90077-6. PubMed PMID: 6178481.

53. Maddaloni G, Bertero A, Pratelli M, Barsotti N, Boonstra A, Giorgi A, Migliarini S, Pasqualetti M. Development of Serotonergic Fibers in the Post-Natal Mouse Brain. Fron Cell Neurosci. 2017;11:202. Epub 20170714. doi: 10.3389/fncel.2017.00202. PubMed PMID: 28769763; PMCID: PMC5509955.

54. Vahid-Ansari F, Albert PR. Rewiring of the Serotonin System in Major Depression. Front Psychiatry. 2021;12:802581. Epub 20211216. doi: 10.3389/fpsyt.2021.802581. PubMed PMID: 34975594; PMCID: PMC8716791.

55. Rivera PD, Hanamsagar R, Kan MJ, Tran PK, Stewart D, Jo YC, Gunn M, Bilbo SD. Removal of microglial-specific MyD88 signaling alters dentate gyrus doublecortin and enhances opioid addiction-like behaviors. Brain Behav Immun. 2019;76:104–15. Epub 20181115. doi: 10.1016/j.bbi.2018.11.010. PubMed PMID: 30447281; PMCID: PMC6348129.

56. Deng F, Wan J, Li G, Dong H, Xia X, Wang Y, Li X, Zhuang C, Zheng Y, Liu L, Yan Y, Feng J, Zhao Y, Xie H, Li Y. Improved green and red GRAB sensors for monitoring spatiotemporal serotonin release in vivo. Nat Methods. 2024;21(4):692–702. Epub 20240305. doi: 10.1038/s41592-024-02188-8. PubMed PMID: 38443508; PMCID: PMC11377854.

57. Lv Z, Li L, Li Y, Zhang L, Guo X, Huang C, Hou W, Qu Y, Liu L, Li Y, He Z, Tai F. Involvement of Serotonergic Projections from the Dorsal Raphe to the Medial Preoptic Area in the Regulation of the Pup-Directed Paternal Response of Male Mandarin Voles. Int J Mol Sci. 2023;24(14). Epub 20230718. doi: 10.3390/ijms241411605. PubMed PMID: 37511364; PMCID: PMC10380723.

58. Okada Y, Hosoi N, Matsuzaki Y, Fukai Y, Hiraga A, Nakai J, Nitta K, Shinohara Y, Konno A, Hirai H. Development of microglia-targeting adeno-associated viral vectors as tools to study microglial behavior in vivo. Commun Biol. 2022;5(1):1224. Epub 20221111. doi: 10.1038/s42003-022-04200-3. PubMed PMID: 36369525; PMCID: PMC9652230.

59. Aoki R, Konno A, Hosoi N, Kawabata H, Hirai H. AAV vectors for specific and efficient gene expression in microglia. Cell Rep Methods. 2025;5(8):101116. Epub 20250730. doi: 10.1016/j.crmeth.2025.101116. PubMed PMID: 40744011; PMCID: PMC12461629.

60. Patton MS, Sheats SH, Siclair AN, Mathur BN. Alcohol potentiates multiple GABAergic inputs to dorsal striatum fast-spiking interneurons. Neuropharmacology. 2023;232:109527. Epub 20230401. doi: 10.1016/j.neuropharm.2023.109527. PubMed PMID: 37011784; PMCID: PMC10122715.

61. Patton MS, Sheats SH, Wulff AB, McKeon PN, VanRyzin JW, Patton MH, Heckman M, Siclair AN, Iffland PH, Mathur BN. Perineuronal Net and Inhibitory Synapse Remodeling on Striatal Fast-spiking Interneurons by Chronic Alcohol Exposure. bioRxiv. 2025. Epub 20250712. doi: 10.1101/2025.07.08.663744. PubMed PMID: 40672273; PMCID: PMC12265726.

62. Devlin BA, Nguyen DM, Ribeiro D, Grullon G, Clark MJ, Finn A, Ceasrine AM, Oxendine S, Deja M, Shah A, Ati S, Schaefer A, Bilbo SD. Excitatory Neuron-Derived Interleukin-34 Controls Cortical Developmental Microglia Function. bioRxiv. 2025. Epub 20250507. doi: 10.1101/2024.05.10.589920. PubMed PMID: 38766127; PMCID: PMC11100801.

63. Berg S, Kutra D, Kroeger T, Straehle CN, Kausler BX, Haubold C, Schiegg M, Ales J, Beier T, Rudy M, Eren K, Cervantes JI, Xu B, Beuttenmueller F, Wolny A, Zhang C, Koethe U, Hamprecht FA, Kreshuk A. ilastik: interactive machine learning for (bio)image analysis. Nat Methods. 2019;16(12):1226–32. Epub 20190930. doi: 10.1038/s41592-019-0582-9. PubMed PMID: 31570887.

64. Yates SC, Groeneboom NE, Coello C, Lichtenthaler SF, Kuhn PH, Demuth HU, Hartlage-Rübsamen M, Roßner S, Leergaard T, Kreshuk A, Puchades MA, Bjaalie JG. QUINT: Workflow for Quantification and Spatial Analysis of Features in Histological Images From Rodent Brain. Front Neuroinform. 2019;13:75. Epub 20191203. doi: 10.3389/fninf.2019.00075. PubMed PMID: 31849633; PMCID: PMC6901597.

65. Groeneboom NE, Yates SC, Puchades MA, Bjaalie JG. Nutil: A Pre- and Post-processing Toolbox for Histological Rodent Brain Section Images. Front Neuroinform. 2020;14:37. Epub 20200821. doi: 10.3389/fninf.2020.00037. PubMed PMID: 32973479; PMCID: PMC7472695.

66. Sanz E, Bean JC, Carey DP, Quintana A, McKnight GS. RiboTag: Ribosomal Tagging Strategy to Analyze Cell-Type-Specific mRNA Expression In Vivo. Curr Protoc Neurosci. 2019;88(1):e77. doi: 10.1002/cpns.77. PubMed PMID: 31216392; PMCID: PMC6615552.

67. Pallares LF, Picard S, Ayroles JF. TM3’seq: A Tagmentation-Mediated 3’ Sequencing Approach for Improving Scalability of RNAseq Experiments. G3 (Bethesda). 2020;10(1):143–50. Epub 20200107. doi: 10.1534/g3.119.400821. PubMed PMID: 31676507; PMCID: PMC6945013.

68. Bordt EA, Block CL, Petrozziello T, Sadri-Vakili G, Smith CJ, Edlow AG, Bilbo SD. Isolation of Microglia from Mouse or Human Tissue. STAR Protoc. 2020;1(1). Epub 20200603. doi: 10.1016/j.xpro.2020.100035. PubMed PMID: 32783030; PMCID: PMC7416840.

69. Bergstrom HC, Lieberman AG, Graybeal C, Lipkin AM, Holmes A. Dorsolateral striatum engagement during reversal learning. Learn Mem. 2020;27(10):418–22. Epub 20200915. doi: 10.1101/lm.051714.120. PubMed PMID: 32934094; PMCID: PMC7497112.

70. Rhodes JS, Best K, Belknap JK, Finn DA, Crabbe JC. Evaluation of a simple model of ethanol drinking to intoxication in C57BL/6J mice. Physiol Behav. 2005;84(1):53–63. doi: 10.1016/j.physbeh.2004.10.007. PubMed PMID: 15642607.

71. Patton MS, Heckman M, Kim C, Mu C, Mathur BN. Compulsive alcohol consumption is regulated by dorsal striatum fast-spiking interneurons. Neuropsychopharmacology. 2021;46(2):351–9. Epub 20200714. doi: 10.1038/s41386-020-0766-0. PubMed PMID: 32663841; PMCID: PMC7852608.

72. Nagai Y, Miyakawa N, Takuwa H, Hori Y, Oyama K, Ji B, Takahashi M, Huang XP, Slocum ST, DiBerto JF, Xiong Y, Urushihata T, Hirabayashi T, Fujimoto A, Mimura K, English JG, Liu J, Inoue KI, Kumata K, Seki C, Ono M, Shimojo M, Zhang MR, Tomita Y, Nakahara J, Suhara T, Takada M, Higuchi M, Jin J, Roth BL, Minamimoto T. Deschloroclozapine, a potent and selective chemogenetic actuator enables rapid neuronal and behavioral modulations in mice and monkeys. Nat Neurosci. 2020;23(9):1157–67. Epub 20200706. doi: 10.1038/s41593-020-0661-3. PubMed PMID: 32632286.

73. Huang KW, Ochandarena NE, Philson AC, Hyun M, Birnbaum JE, Cicconet M, Sabatini BL. Molecular and anatomical organization of the dorsal raphe nucleus. Elife. 2019;8. Epub 20190814. doi: 10.7554/eLife.46464. PubMed PMID: 31411560; PMCID: PMC6726424.

74. Ren J, Friedmann D, Xiong J, Liu CD, Ferguson BR, Weerakkody T, DeLoach KE, Ran C, Pun A, Sun Y, Weissbourd B, Neve RL, Huguenard J, Horowitz MA, Luo L. Anatomically Defined and Functionally Distinct Dorsal Raphe Serotonin Sub-systems. Cell. 2018;175(2):472–87.e20. Epub 20180823. doi: 10.1016/j.cell.2018.07.043. PubMed PMID: 30146164; PMCID: PMC6173627.

75. Deneris E, Gaspar P. Serotonin neuron development: shaping molecular and structural identities. Wiley Interdiscip Rev Dev Biol. 2018;7(1). Epub 20171026. doi: 10.1002/wdev.301. PubMed PMID: 29072810; PMCID: PMC5746461.

76. Belmer A, Klenowski PM, Patkar OL, Bartlett SE. Mapping the connectivity of serotonin transporter immunoreactive axons to excitatory and inhibitory neurochemical synapses in the mouse limbic brain. Brain Struct Funct. 2017;222(3):1297–314. Epub 20160802. doi: 10.1007/s00429-016-1278-x. PubMed PMID: 27485750; PMCID: PMC5368196.

77. Hoops D, Reynolds LM, Restrepo-Lozano JM, Flores C. Dopamine Development in the Mouse Orbital Prefrontal Cortex Is Protracted and Sensitive to Amphetamine in Adolescence. eNeuro. 2018;5(1). Epub 20180110. doi: 10.1523/eneuro.0372-17.2017. PubMed PMID: 29333488; PMCID: PMC5762649.

78. Levitt P, Moore RY. Development of the noradrenergic innervation of neocortex. Brain Res. 1979;162(2):243–59. doi: 10.1016/0006-8993(79)90287-7. PubMed PMID: 761089.

79. Loy R, Moore RY. Ontogeny of the noradrenergic innervation of the rat hippocampal formation. Anat Embryol (Berl). 1979;157(3):243–53. doi: 10.1007/bf00304992. PubMed PMID: 525817.

80. Kolodziejczak M, Béchade C, Gervasi N, Irinopoulou T, Banas SM, Cordier C, Rebsam A, Roumier A, Maroteaux L. Serotonin Modulates Developmental Microglia via 5-HT2B Receptors: Potential Implication during Synaptic Refinement of Retinogeniculate Projections. ACS Chem Neurosci. 2015;6(7):1219–30. Epub 20150417. doi: 10.1021/cn5003489. PubMed PMID: 25857335.

81. Béchade C, D’Andrea I, Etienne F, Verdonk F, Moutkine I, Banas SM, Kolodziejczak M, Diaz SL, Parkhurst CN, Gan WB, Maroteaux L, Roumier A. The serotonin 2B receptor is required in neonatal microglia to limit neuroinflammation and sickness behavior in adulthood. Glia. 2021;69(3):638–54. Epub 20201023. doi: 10.1002/glia.23918. PubMed PMID: 33095507.

82. Albertini G, D’Andrea I, Druart M, Béchade C, Nieves-Rivera N, Etienne F, Le Magueresse C, Rebsam A, Heck N, Maroteaux L, Roumier A. Serotonin sensing by microglia conditions the proper development of neuronal circuits and of social and adaptive skills. Mol Psychiatry. 2023;28(6):2328–42. Epub 20230522. doi: 10.1038/s41380-023-02048-5. PubMed PMID: 37217677.

83. Krabbe G, Matyash V, Pannasch U, Mamer L, Boddeke HW, Kettenmann H. Activation of serotonin receptors promotes microglial injury-induced motility but attenuates phagocytic activity. Brain Behav Immun. 2012;26(3):419–28. Epub 20111217. doi: 10.1016/j.bbi.2011.12.002. PubMed PMID: 22198120.

84. Glebov K, Löchner M, Jabs R, Lau T, Merkel O, Schloss P, Steinhäuser C, Walter J. Serotonin stimulates secretion of exosomes from microglia cells. Glia. 2015;63(4):626–34. Epub 20141201. doi: 10.1002/glia.22772. PubMed PMID: 25451814.

85. Zengel J, Wang YX, Seo JW, Ning K, Hamilton JN, Wu B, Raie M, Holbrook C, Su S, Clements DR, Pillay S, Puschnik AS, Winslow MM, Idoyaga J, Nagamine CM, Sun Y, Mahajan VB, Ferrara KW, Blau HM, Carette JE. Hardwiring tissue-specific AAV transduction in mice through engineered receptor expression. Nat Methods. 2023;20(7):1070–81. Epub 20230608. doi: 10.1038/s41592-023-01896-x. PubMed PMID: 37291262; PMCID: PMC10333121.

86. Maes ME, Colombo G, Schulz R, Siegert S. Targeting microglia with lentivirus and AAV: Recent advances and remaining challenges. Neurosci Lett. 2019;707:134310. Epub 20190531. doi: 10.1016/j.neulet.2019.134310. PubMed PMID: 31158432; PMCID: PMC6734419.

87. Stamataki M, Rissiek B, Magnus T, Körbelin J. Microglia targeting by adeno-associated viral vectors. Front Immunol. 2024;15:1425892. Epub 20240705. doi: 10.3389/fimmu.2024.1425892. PubMed PMID: 39035004; PMCID: PMC11257843.

88. Ren J, Isakova A, Friedmann D, Zeng J, Grutzner SM, Pun A, Zhao GQ, Kolluru SS, Wang R, Lin R, Li P, Li A, Raymond JL, Luo Q, Luo M, Quake SR, Luo L. Single-cell transcriptomes and whole-brain projections of serotonin neurons in the mouse dorsal and median raphe nuclei. Elife. 2019;8. Epub 20191024. doi: 10.7554/eLife.49424. PubMed PMID: 31647409; PMCID: PMC6812963.

89. Okaty BW, Sturrock N, Escobedo Lozoya Y, Chang Y, Senft RA, Lyon KA, Alekseyenko OV, Dymecki SM. A single-cell transcriptomic and anatomic atlas of mouse dorsal raphe Pet1 neurons. Elife. 2020;9. Epub 20200622. doi: 10.7554/eLife.55523. PubMed PMID: 32568072; PMCID: PMC7308082.

90. Brummelte S, Mc Glanaghy E, Bonnin A, Oberlander TF. Developmental changes in serotonin signaling: Implications for early brain function, behavior and adaptation. Neuroscience. 2017;342:212–31. Epub 20160222. doi: 10.1016/j.neuroscience.2016.02.037. PubMed PMID: 26905950; PMCID: PMC5310545.

91. van Kleef ES, Gaspar P, Bonnin A. Insights into the complex influence of 5-HT signaling on thalamocortical axonal system development. Eur J Neurosci. 2012;35(10):1563–72. doi: 10.1111/j.1460-9568.2012.8096.x. PubMed PMID: 22607002; PMCID: PMC3359868.

92. Miceli S, Negwer M, van Eijs F, Kalkhoven C, van Lierop I, Homberg J, Schubert D. High serotonin levels during brain development alter the structural input-output connectivity of neural networks in the rat somatosensory layer IV. Front Cell Neurosci. 2013;7:88. Epub 20130607. doi: 10.3389/fncel.2013.00088. PubMed PMID: 23761736; PMCID: PMC3675331.

93. Lebrand C, Cases O, Adelbrecht C, Doye A, Alvarez C, El Mestikawy S, Seif I, Gaspar P. Transient uptake and storage of serotonin in developing thalamic neurons. Neuron. 1996;17(5):823–35. doi: 10.1016/s0896-6273(00)80215-9. PubMed PMID: 8938116.

94. Riccio O, Potter G, Walzer C, Vallet P, Szabó G, Vutskits L, Kiss JZ, Dayer AG. Excess of serotonin affects embryonic interneuron migration through activation of the serotonin receptor 6. Mol Psychiatry. 2009;14(3):280–90. Epub 20080729. doi: 10.1038/mp.2008.89. PubMed PMID: 18663366.

95. Corbit LH, Muir JL, Balleine BW. The role of the nucleus accumbens in instrumental conditioning: Evidence of a functional dissociation between accumbens core and shell. J Neurosci. 2001;21(9):3251–60. doi: 10.1523/jneurosci.21-09-03251.2001. PubMed PMID: 11312310; PMCID: PMC6762583.

96. Patton MS, Lodge DJ, Morilak DA, Girotti M. Ketamine Corrects Stress-Induced Cognitive Dysfunction through JAK2/STAT3 Signaling in the Orbitofrontal Cortex. Neuropsychopharmacology. 2017;42(6):1220–30. Epub 20161017. doi: 10.1038/npp.2016.236. PubMed PMID: 27748739; PMCID: PMC5437880.

97. Clarke HF, Dalley JW, Crofts HS, Robbins TW, Roberts AC. Cognitive inflexibility after prefrontal serotonin depletion. Science. 2004;304(5672):878–80. doi: 10.1126/science.1094987. PubMed PMID: 15131308.

98. Clarke HF, Walker SC, Crofts HS, Dalley JW, Robbins TW, Roberts AC. Prefrontal serotonin depletion affects reversal learning but not attentional set shifting. J Neurosci. 2005;25(2):532–8. doi: 10.1523/jneurosci.3690-04.2005. PubMed PMID: 15647499; PMCID: PMC6725478.

99. Clarke HF, Robbins TW, Roberts AC. Lesions of the medial striatum in monkeys produce perseverative impairments during reversal learning similar to those produced by lesions of the orbitofrontal cortex. J Neurosci. 2008;28(43):10972–82. doi: 10.1523/jneurosci.1521-08.2008. PubMed PMID: 18945905; PMCID: PMC3981993.

100. Matias S, Lottem E, Dugué GP, Mainen ZF. Activity patterns of serotonin neurons underlying cognitive flexibility. Elife. 2017;6. Epub 20170321. doi: 10.7554/eLife.20552. PubMed PMID: 28322190; PMCID: PMC5360447.

101. Gourley SL, Olevska A, Zimmermann KS, Ressler KJ, Dileone RJ, Taylor JR. The orbitofrontal cortex regulates outcome-based decision-making via the lateral striatum. Eur J Neurosci. 2013;38(3):2382–8. Epub 20130508. doi: 10.1111/ejn.12239. PubMed PMID: 23651226; PMCID: PMC3864662.

102. Panayi MC, Killcross S. Functional heterogeneity within the rodent lateral orbitofrontal cortex dissociates outcome devaluation and reversal learning deficits. Elife. 2018;7. Epub 20180725. doi: 10.7554/eLife.37357. PubMed PMID: 30044220; PMCID: PMC6101941.

103. Gourley SL, Lee AS, Howell JL, Pittenger C, Taylor JR. Dissociable regulation of instrumental action within mouse prefrontal cortex. Eur J Neurosci. 2010;32(10):1726–34. Epub 20101012. doi: 10.1111/j.1460-9568.2010.07438.x. PubMed PMID: 21044173; PMCID: PMC3058746.

104. Izquierdo A, Brigman JL, Radke AK, Rudebeck PH, Holmes A. The neural basis of reversal learning: An updated perspective. Neuroscience. 2017;345:12–26. Epub 20160312. doi: 10.1016/j.neuroscience.2016.03.021. PubMed PMID: 26979052; PMCID: PMC5018909.

105. Corbit LH, Balleine BW. The general and outcome-specific forms of Pavlovian-instrumental transfer are differentially mediated by the nucleus accumbens core and shell. J Neurosci. 2011;31(33):11786–94. doi: 10.1523/jneurosci.2711-11.2011. PubMed PMID: 21849539; PMCID: PMC3208020.

106. Fritz M, El Rawas R, Klement S, Kummer K, Mayr MJ, Eggart V, Salti A, Bardo MT, Saria A, Zernig G. Differential effects of accumbens core vs. shell lesions in a rat concurrent conditioned place preference paradigm for cocaine vs. social interaction. PLoS One. 2011;6(10):e26761. Epub 20111026. doi: 10.1371/journal.pone.0026761. PubMed PMID: 22046347; PMCID: PMC3202564.

107. Nennig SE, Fulenwider HD, Chimberoff SH, Smith BM, Eskew JE, Sequeira MK, Karlsson C, Liang C, Chen JF, Heilig M, Schank JR. Selective Lesioning of Nuclear Factor-κB Activated Cells in the Nucleus Accumbens Shell Attenuates Alcohol Place Preference. Neuropsychopharmacology. 2018;43(5):1032–40. Epub 20170913. doi: 10.1038/npp.2017.214. PubMed PMID: 28901327; PMCID: PMC5854796.

108. Carr GD, White NM. Conditioned place preference from intra-accumbens but not intra-caudate amphetamine injections. Life Sci. 1983;33(25):2551–7. doi: 10.1016/0024-3205(83)90165-0. PubMed PMID: 6645814.

109. Li Y, Zhong W, Wang D, Feng Q, Liu Z, Zhou J, Jia C, Hu F, Zeng J, Guo Q, Fu L, Luo M. Serotonin neurons in the dorsal raphe nucleus encode reward signals. Nat Commun. 2016;7:10503. Epub 20160128. doi: 10.1038/ncomms10503. PubMed PMID: 26818705; PMCID: PMC4738365.

110. Ranade SP, Mainen ZF. Transient firing of dorsal raphe neurons encodes diverse and specific sensory, motor, and reward events. J Neurophysiol. 2009;102(5):3026–37. Epub 20090826. doi: 10.1152/jn.00507.2009. PubMed PMID: 19710375.

111. Cohen JY, Amoroso MW, Uchida N. Serotonergic neurons signal reward and punishment on multiple timescales. Elife. 2015;4. Epub 20150225. doi: 10.7554/eLife.06346. PubMed PMID: 25714923; PMCID: PMC4389268.

112. Liu Z, Zhou J, Li Y, Hu F, Lu Y, Ma M, Feng Q, Zhang JE, Wang D, Zeng J, Bao J, Kim JY, Chen ZF, El Mestikawy S, Luo M. Dorsal raphe neurons signal reward through 5-HT and glutamate. Neuron. 2014;81(6):1360–74. doi: 10.1016/j.neuron.2014.02.010. PubMed PMID: 24656254; PMCID: PMC4411946.

113. Purohit K, Parekh PK, Kern J, Logan RW, Liu Z, Huang Y, McClung CA, Crabbe JC, Ozburn AR. Pharmacogenetic Manipulation of the Nucleus Accumbens Alters Binge-Like Alcohol Drinking in Mice. Alcohol Clin Exp Res. 2018;42(5):879–88. Epub 20180418. doi: 10.1111/acer.13626. PubMed PMID: 29668112; PMCID: PMC6034712.

114. Townsley KG, Borrego MB, Ozburn AR. Effects of chemogenetic manipulation of the nucleus accumbens core in male C57BL/6J mice. Alcohol. 2021;91:21–7. Epub 20201104. doi: 10.1016/j.alcohol.2020.10.005. PubMed PMID: 33160072; PMCID: PMC8675149.

115. Kwako LE, Koob GF. Neuroclinical Framework for the Role of Stress in Addiction. Chronic Stress (Thousand Oaks). 2017;1. Epub 20170410. doi: 10.1177/2470547017698140. PubMed PMID: 28653044; PMCID: PMC5482275.

116. Pickard GE, Weber ET, Scott PA, Riberdy AF, Rea MA. 5HT1B receptor agonists inhibit light-induced phase shifts of behavioral circadian rhythms and expression of the immediate-early gene c-fos in the suprachiasmatic nucleus. J Neurosci. 1996;16(24):8208–20. doi: 10.1523/jneurosci.16-24-08208.1996. PubMed PMID: 8987845; PMCID: PMC6579213.

117. Pickard GE, Smith BN, Belenky M, Rea MA, Dudek FE, Sollars PJ. 5-HT1B receptor-mediated presynaptic inhibition of retinal input to the suprachiasmatic nucleus. J Neurosci. 1999;19(10):4034–45. doi: 10.1523/jneurosci.19-10-04034.1999. PubMed PMID: 10234032; PMCID: PMC6782735.

118. Edlow AG. Maternal Metabolic Disease and Offspring Neurodevelopment-An Evolving Public Health Crisis. JAMA Netw Open. 2021;4(10):e2129674. Epub 20211001. doi: 10.1001/jamanetworkopen.2021.29674. PubMed PMID: 34648016.

119. Eleftheriades A, Koulouraki S, Belegrinos A, Eleftheriades M, Pervanidou P. Maternal Obesity and Neurodevelopment of the Offspring. Nutrients. 2025;17(5). Epub 20250302. doi: 10.3390/nu17050891. PubMed PMID: 40077761; PMCID: PMC11901708.

120. Schmitt LO, Faraco G, Stivanin TS, Gaspar JM. Maternal Obesity in Pregnancy: Risk Factor for Neurodevelopmental Outcomes in Offspring. J Neurochem. 2025;169(12):e70333. doi: 10.1111/jnc.70333. PubMed PMID: 41452351; PMCID: PMC12742557.

121. Windham GC, Anderson M, Lyall K, Daniels JL, Kral TVE, Croen LA, Levy SE, Bradley CB, Cordero C, Young L, Schieve LA. Maternal Pre-pregnancy Body Mass Index and Gestational Weight Gain in Relation to Autism Spectrum Disorder and other Developmental Disorders in Offspring. Autism Res. 2019;12(2):316–27. Epub 20181221. doi: 10.1002/aur.2057. PubMed PMID: 30575327; PMCID: PMC7778460.

122. Flegal KM, Carroll MD, Kit BK, Ogden CL. Prevalence of obesity and trends in the distribution of body mass index among US adults, 1999-2010. Jama. 2012;307(5):491–7. Epub 20120117. doi: 10.1001/jama.2012.39. PubMed PMID: 22253363.

123. Stüber TN, Künzel EC, Zollner U, Rehn M, Wöckel A, Hönig A. Prevalence and Associated Risk Factors for Obesity During Pregnancy Over Time. Geburtshilfe Frauenheilkd. 2015;75(9):923–8. doi: 10.1055/s-0035-1557868. PubMed PMID: 26500368; PMCID: PMC4596699.

124. Shaw KA, Williams S, Patrick ME, Valencia-Prado M, Durkin MS, Howerton EM, Ladd-Acosta CM, Pas ET, Bakian AV, Bartholomew P, Nieves-Muñoz N, Sidwell K, Alford A, Bilder DA, DiRienzo M, Fitzgerald RT, Furnier SM, Hudson AE, Pokoski OM, Shea L, Tinker SC, Warren Z, Zahorodny W, Agosto-Rosa H, Anbar J, Chavez KY, Esler A, Forkner A, Grzybowski A, Agib AH, Hallas L, Lopez M, Magaña S, Nguyen RHN, Parker J, Pierce K, Protho T, Torres H, Vanegas SB, Vehorn A, Zhang M, Andrews J, Greer F, Hall-Lande J, McArthur D, Mitamura M, Montes AJ, Pettygrove S, Shenouda J, Skowyra C, Washington A, Maenner MJ. Prevalence and Early Identification of Autism Spectrum Disorder Among Children Aged 4 and 8 Years - Autism and Developmental Disabilities Monitoring Network, 16 Sites, United States, 2022. MMWR Surveill Summ. 2025;74(2):1–22. Epub 20250417. doi: 10.15585/mmwr.ss7402a1. PubMed PMID: 40232988; PMCID: PMC12011386.

125. Grosvenor LP, Croen LA, Lynch FL, Marafino BJ, Maye M, Penfold RB, Simon GE, Ames JL. Autism Diagnosis Among US Children and Adults, 2011-2022. JAMA Netw Open. 2024;7(10):e2442218. Epub 20241001. doi: 10.1001/jamanetworkopen.2024.42218. PubMed PMID: 39476234; PMCID: PMC11525601.

126. Xu G, Strathearn L, Liu B, Yang B, Bao W. Twenty-Year Trends in Diagnosed Attention-Deficit/Hyperactivity Disorder Among US Children and Adolescents, 1997-2016. JAMA Netw Open. 2018;1(4):e181471. Epub 20180803. doi: 10.1001/jamanetworkopen.2018.1471. PubMed PMID: 30646132; PMCID: PMC6324288.

127. Danielson ML, Claussen AH, Bitsko RH, Katz SM, Newsome K, Blumberg SJ, Kogan MD, Ghandour R. ADHD Prevalence Among U.S. Children and Adolescents in 2022: Diagnosis, Severity, Co-Occurring Disorders, and Treatment. J Clin Child Adolesc Psychol. 2024;53(3):343–60. Epub 20240522. doi: 10.1080/15374416.2024.2335625. PubMed PMID: 38778436; PMCID: PMC11334226.

128. Boyle CA, Boulet S, Schieve LA, Cohen RA, Blumberg SJ, Yeargin-Allsopp M, Visser S, Kogan MD. Trends in the prevalence of developmental disabilities in US children, 1997-2008. Pediatrics. 2011;127(6):1034–42. Epub 20110523. doi: 10.1542/peds.2010-2989. PubMed PMID: 21606152.

129. Krakowiak P, Walker CK, Bremer AA, Baker AS, Ozonoff S, Hansen RL, Hertz-Picciotto I. Maternal metabolic conditions and risk for autism and other neurodevelopmental disorders. Pediatrics. 2012;129(5):e1121–8. Epub 20120409. doi: 10.1542/peds.2011-2583. PubMed PMID: 22492772; PMCID: PMC3340592.

130. Schaefer CA, Brown AS, Wyatt RJ, Kline J, Begg MD, Bresnahan MA, Susser ES. Maternal prepregnant body mass and risk of schizophrenia in adult offspring. Schizophr Bull. 2000;26(2):275–86. doi: 10.1093/oxfordjournals.schbul.a033452. PubMed PMID: 10885630.

131. Rodriguez A. Maternal pre-pregnancy obesity and risk for inattention and negative emotionality in children. J Child Psychol Psychiatry. 2010;51(2):134–43. Epub 20090806. doi: 10.1111/j.1469-7610.2009.02133.x. PubMed PMID: 19674195.

132. Urbonaite G, Knyzeliene A, Bunn FS, Smalskys A, Neniskyte U. The impact of maternal high-fat diet on offspring neurodevelopment. Front Neurosci. 2022;16:909762. Epub 20220722. doi: 10.3389/fnins.2022.909762. PubMed PMID: 35937892; PMCID: PMC9354026.

133. Tremblay M, Lowery RL, Majewska AK. Microglial interactions with synapses are modulated by visual experience. PLoS Biol. 2010;8(11):e1000527. Epub 20101102. doi: 10.1371/journal.pbio.1000527. PubMed PMID: 21072242; PMCID: PMC2970556.

134. Miyamoto A, Wake H, Ishikawa AW, Eto K, Shibata K, Murakoshi H, Koizumi S, Moorhouse AJ, Yoshimura Y, Nabekura J. Microglia contact induces synapse formation in developing somatosensory cortex. Nat Commun. 2016;7:12540. Epub 20160825. doi: 10.1038/ncomms12540. PubMed PMID: 27558646; PMCID: PMC5007295.

135. Mathur BN, Capik NA, Alvarez VA, Lovinger DM. Serotonin induces long-term depression at corticostriatal synapses. J Neurosci. 2011;31(20):7402–11. doi: 10.1523/jneurosci.6250-10.2011. PubMed PMID: 21593324; PMCID: PMC3113491.

136. Browne CJ, Ji X, Higgins GA, Fletcher PJ, Harvey-Lewis C. Pharmacological Modulation of 5-HT(2C) Receptor Activity Produces Bidirectional Changes in Locomotor Activity, Responding for a Conditioned Reinforcer, and Mesolimbic DA Release in C57BL/6 Mice. Neuropsychopharmacology. 2017;42(11):2178–87. Epub 20170613. doi: 10.1038/npp.2017.124. PubMed PMID: 28720903; PMCID: PMC5603805.

137. Navailles S, De Deurwaerdère P, Porras G, Spampinato U. In vivo evidence that 5-HT2C receptor antagonist but not agonist modulates cocaine-induced dopamine outflow in the rat nucleus accumbens and striatum. Neuropsychopharmacology. 2004;29(2):319–26. doi: 10.1038/sj.npp.1300329. PubMed PMID: 14560323.

138. Heifets BD, Salgado JS, Taylor MD, Hoerbelt P, Cardozo Pinto DF, Steinberg EE, Walsh JJ, Sze JY, Malenka RC. Distinct neural mechanisms for the prosocial and rewarding properties of MDMA. Sci Transl Med. 2019;11(522). doi: 10.1126/scitranslmed.aaw6435. PubMed PMID: 31826983; PMCID: PMC7123941.

139. Schwarz JM, Sholar PW, Bilbo SD. Sex differences in microglial colonization of the developing rat brain. J Neurochem. 2012;120(6):948–63. Epub 20120209. doi: 10.1111/j.1471-4159.2011.07630.x. PubMed PMID: 22182318; PMCID: PMC3296888.

140. Bordt EA, Ceasrine AM, Bilbo SD. Microglia and sexual differentiation of the developing brain: A focus on ontogeny and intrinsic factors. Glia. 2020;68(6):1085–99. Epub 20191119. doi: 10.1002/glia.23753. PubMed PMID: 31743527; PMCID: PMC7148120.

141. Klein SL, Flanagan KL. Sex differences in immune responses. Nat Rev Immunol. 2016;16(10):626–38. Epub 20160822. doi: 10.1038/nri.2016.90. PubMed PMID: 27546235.

142. Gal-Oz ST, Shay T. Immune Sexual Dimorphism: Connecting the Dots. Physiology (Bethesda). 2022;37(2):55–68. Epub 20210913. doi: 10.1152/physiol.00006.2021. PubMed PMID: 34514870; PMCID: PMC8873034.

