## Supplementary figures and images for "Maternal high-fat diet drives sex-specific microglia remodeling of serotonergic reward circuitry"

### Extended Data Figure 1

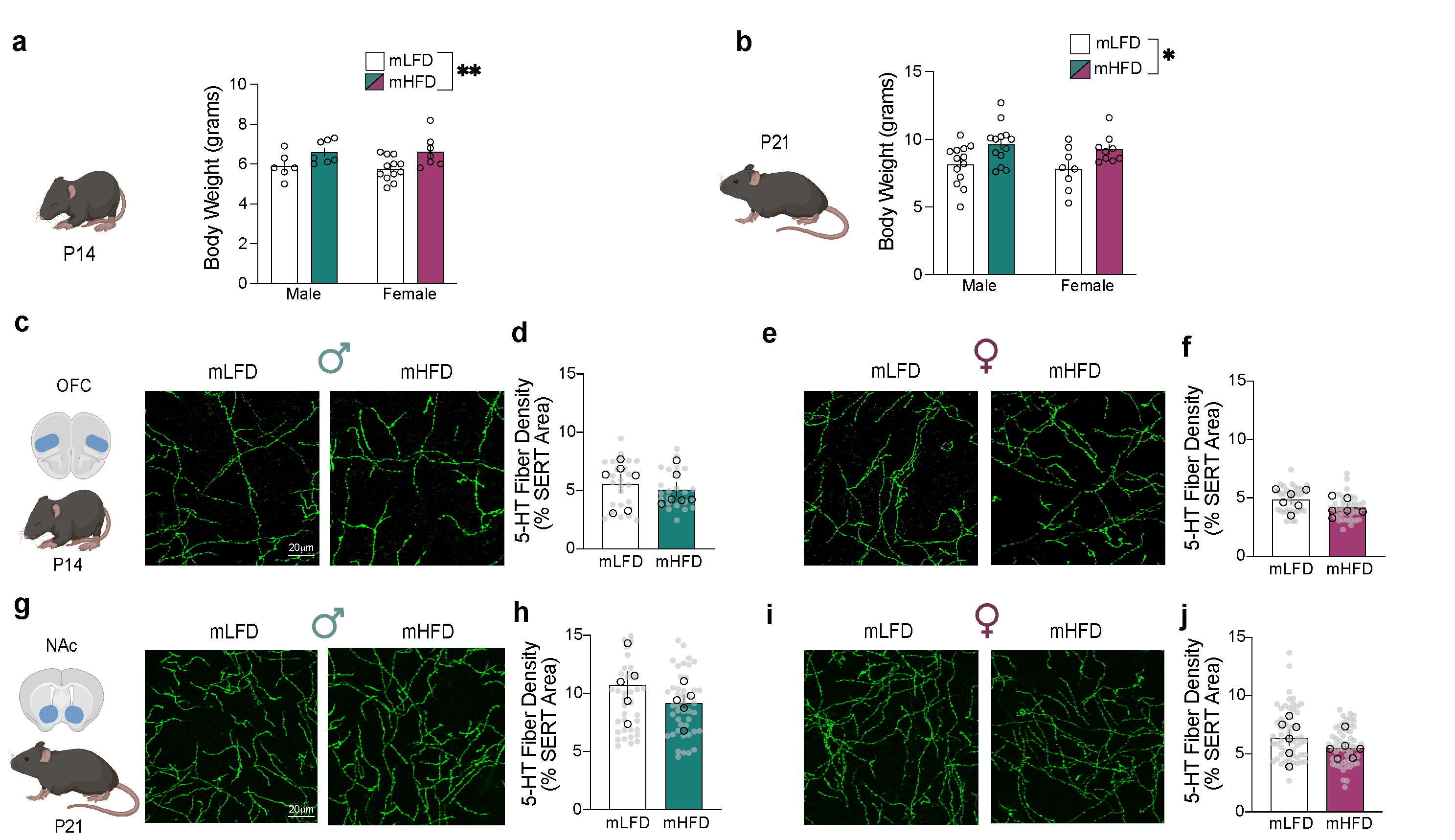

### Extended Data Figure 2

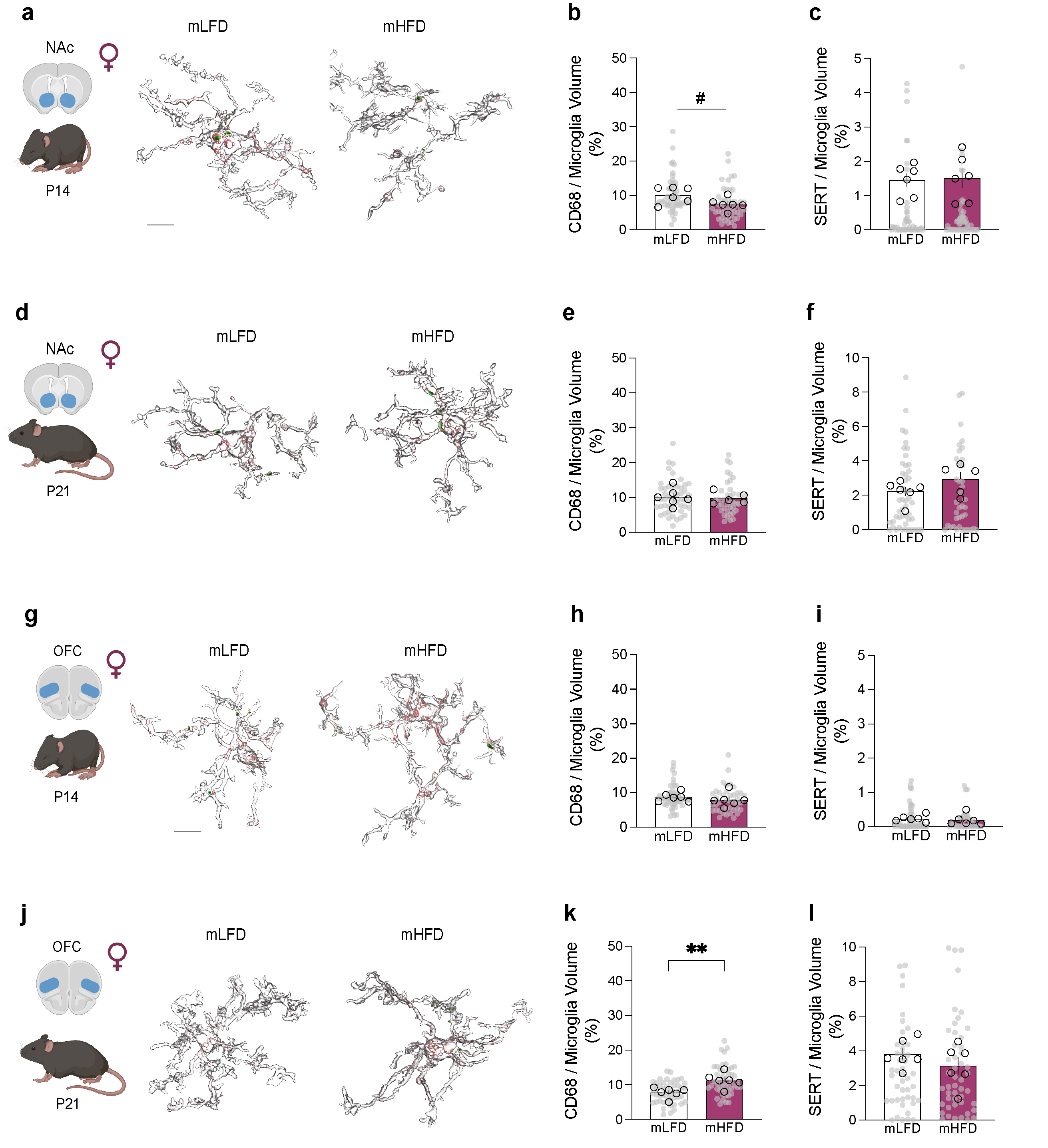

### Extended Data Figure 3

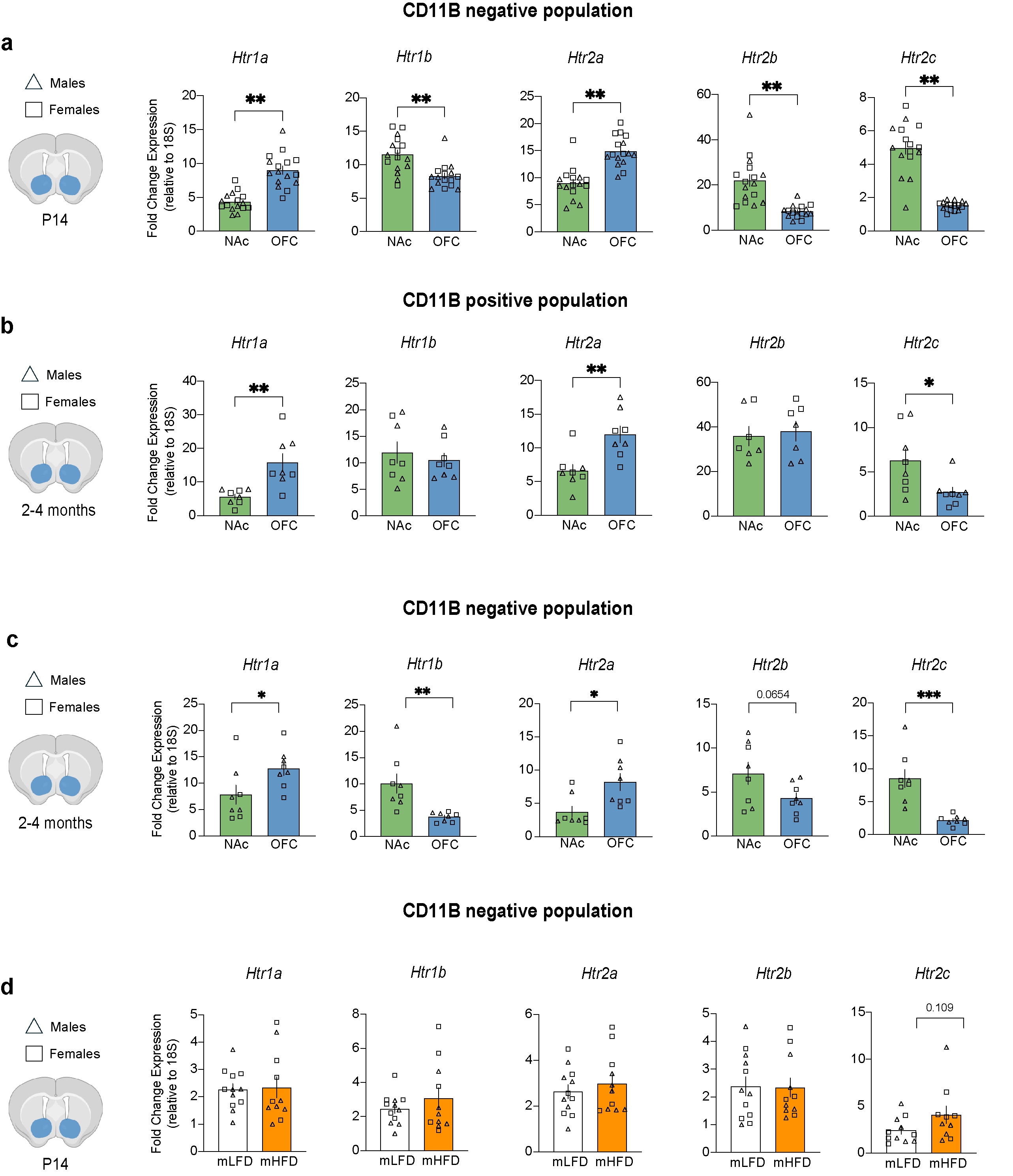

### Extended Data Figure 4

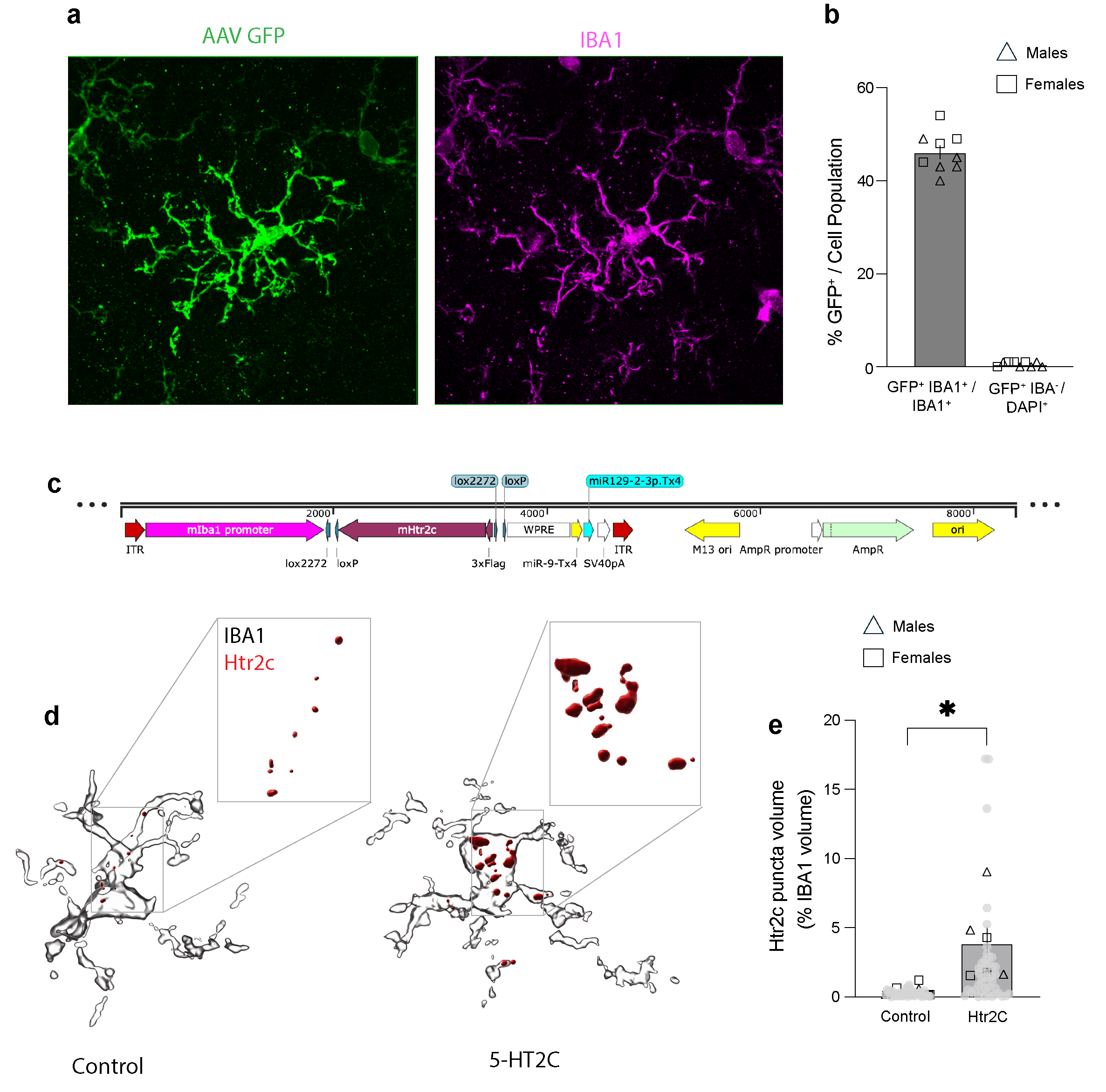

### Extended Data Figure 5

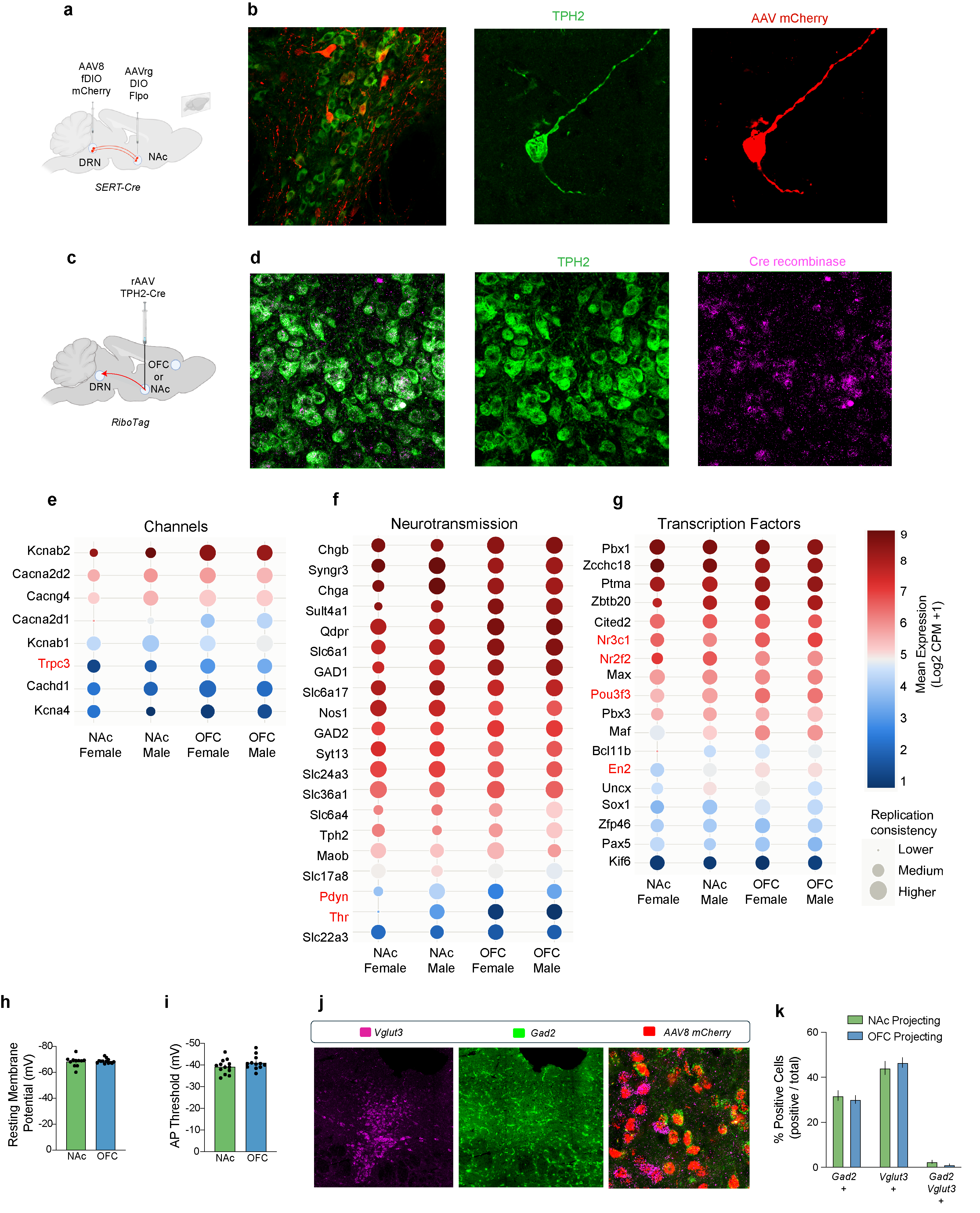

### Extended Data Figure 6

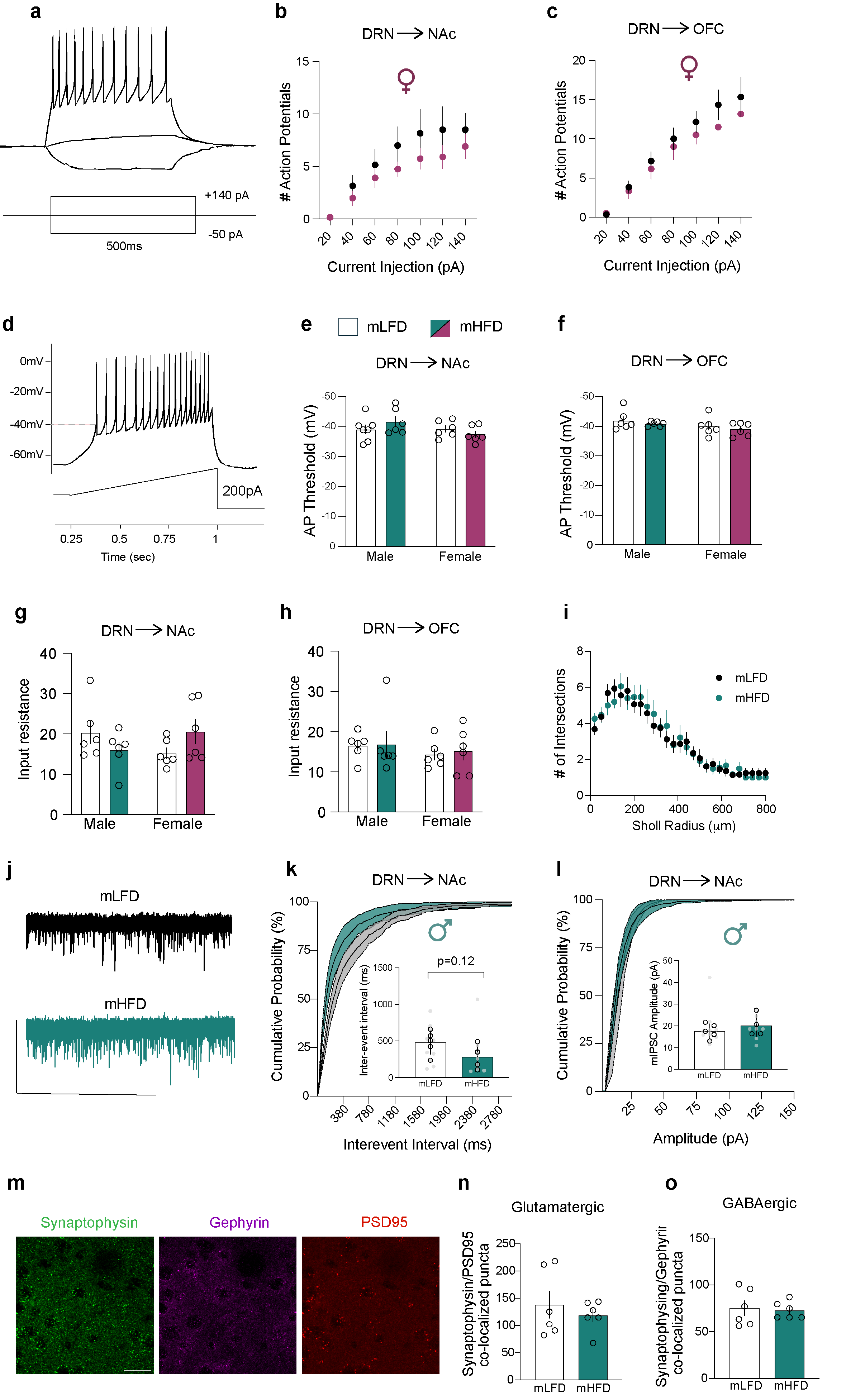

### Extended Data Figure 7

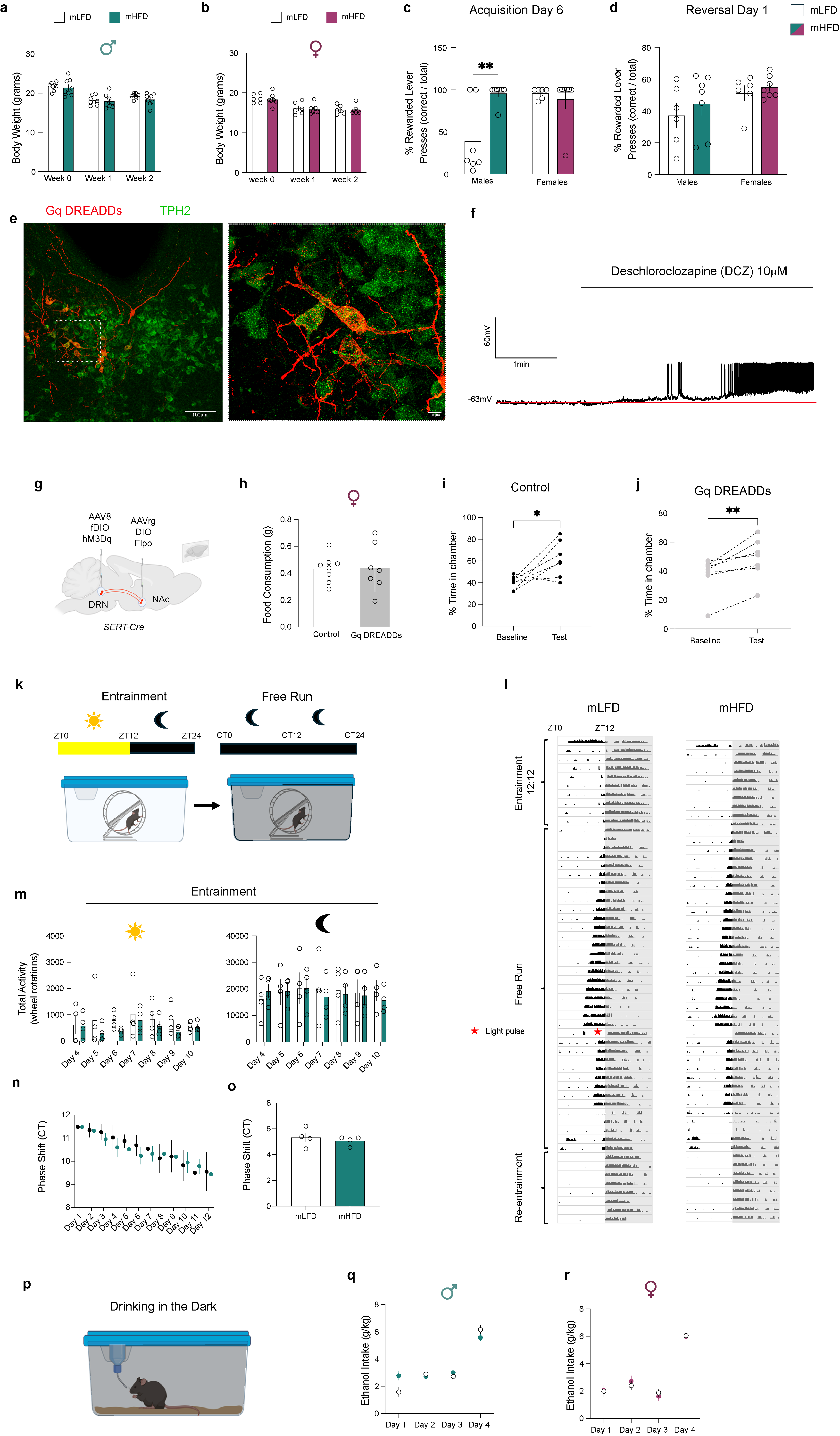
